# Osteoblastic PLEKHO1 contributes to joint inflammation in rheumatoid arthritis

**DOI:** 10.1101/380303

**Authors:** Xiaojuan He, Jin Liu, Chao Liang, Shaikh Atik Badshah, Kang Zheng, Lei Dang, Baosheng Guo, Defang Li, Cheng Lu, Qingqing Guo, Danping Fan, Yanqin Bian, Hui Feng, Lianbo Xiao, Xiaohua Pan, Cheng Xiao, BaoTing Zhang, Ge Zhang, Aiping Lu

## Abstract

Osteoblasts participating in the inflammation regulation gradually obtain concerns. However, its role in joint inflammation of rheumatoid arthritis (RA) is largely unknown. Pleckstrin homology domain-containing family O member 1 (PLEKHO1) was previously identified as a negative regulator of osteogenic lineage activity. Here we demonstrated that PLEKHO1 was highly expressed in osteoblasts of articular specimens from RA patients and inflammatory arthritis mice. Genetic deletion of osteoblastic Plekho1 ameliorated joint inflammation in mice with collagen-induced arthritis (CIA) and K/BxN serum-transfer arthritis (STA), whereas overexpressing Plekho1 only within osteoblasts in CIA and STA mice demonstrated exacerbated local inflammation. Further *in vitro* studies indicated that PLEKHO1 was required for TRAF2-mediated RIP1 ubiquitination to activate NF-kB for inducing inflammatory cytokines production in osteoblasts. Moreover, osteoblastic PLEKHO1 inhibition improved joint inflammation and attenuated bone formation reduction in CIA mice and non-human primate arthritis model. These data strongly suggest that highly expressed PLEKHO1 in osteoblast mediates joint inflammation in RA. Targeting osteoblastic PLEKHO1 may exert dual therapeutic action of alleviating joint inflammation and promoting bone formation in RA.

## Introduction

Rheumatoid arthritis (RA) is characterized by chronic inflammation and progressive bone destruction[1, 2]. Osteoblasts are well-known cell types to modulate bone formation. Interestingly, recent increasing line of evidence suggests that osteoblasts could produce proinflammatory molecules in response to bacterial challenge and contribute to inflammation through the recruitment of leukocytes to the sites of infection during bone diseases such as osteomyelitis[3, 4]. However, whether osteoblasts contribute to local joint inflammation in RA progression and which molecule event is involved in is largely unknown.

Pleckstrin homology domain-containing family O member 1 (PLEKHO1, also known as casein kinase-2 interacting protein-1 (CKIP-1)) is a scaffold protein mediating the interactions with various proteins in multiple signaling pathways, including the osteogenic BMP (bone morphogenetic protein) signaling pathway and the inflammatory PI3K (phosphoinositide 3-kinase)-Akt pathways[5, 6]. Accordingly, the *Plekho1* systemic knock out mice showed higher bone mass than their wide-type controls, indicating that PLEKHO1 may play a role in regulating bone formation[7]. In our previous studies, we further demonstrated that loss of PLEKHO1 in osteoblasts alleviated the age-related bone formation reduction, and osteoblast-targeted Plekho1 siRNA inhibition could promote bone formation in aging and osteoporotic rodents[8, 9]. On the other hand, several recent studies have reported that PLEKHO1 was also involved in cytokine signaling in response to inflammation[10, 11]. However, it remains unclear whether PLEKHO1 contributes to the local joint inflammation and reduction of bone formation in RA pathogenesis.

In this study, we found PLEKHO1 was highly expressed in osteoblasts of joint bone specimens from RA patients and collagen-induced arthritis (CIA) mice. By genetic approach, we found that deletion of osteoblastic *Plekho1* resulted in remarkable amelioration of local inflammation in CIA mice and K/BxN serum-transfer arthritis (STA) mice, whereas genetic overexpression of *Plekho1* only in osteoblasts of CIA mice and STA mice dramatically exacerbated joint inflammation. More importantly, we elucidated that PLEKHO1 was required for TRAF2-mediated RIP1 ubiquitination to activate nuclear factor-k B (NF-kB) for inducing inflammatory cytokines production in osteoblasts. Finally, we also found early inhibition of osteoblastic PLEKHO1 improved joint inflammation and promoted bone formation in CIA mice and the non-human primate arthritis model.

Together, this study uncovers a previously unrecognized role of osteoblastic PLEKHO1 in joint inflammation regulation during RA, and targeting osteoblastic PLEKHO1 could serve as a new pharmacological target with dual action of inflammation inhibition and bone formation augmentation in RA.

## Results

### Osteoblastic PLEKHO1 is upregulated in RA patients and CIA mice

To assess whether PLEKHO1 has a role in RA pathology, we firstly investigated the expression of PLEKHO1 *in vivo.* We collected the bone samples from the knee joint of fifteen RA patients and eight severe trauma (TM) patients underwent knee joint replacement surgery. Indeed, we observed high levels of PLEKHO1 in knee joint bone tissues from RA patients when compared to those from TM controls (**Fig. 1a, 1b**). Further, we detected the levels of PLEKHO1 in osteoblasts by immunofluorescence staining. We used anti-osteocalcin antibody to identify osteoblasts. The results showed an increase of colocalization of PLEKHO1 positive with osteocalcin (Ocn) positive in knee joint bone tissues from RA patients (**Fig. 1c**). Quantitatively, the percentage of cells co-expressing PLEKHO1 and Ocn in Ocn positive (Ocn+) cells of bone tissues from knee joint was also remarkably higher in RA patients compared to TM patients (**Fig. 1d**).

**Figure 1.**
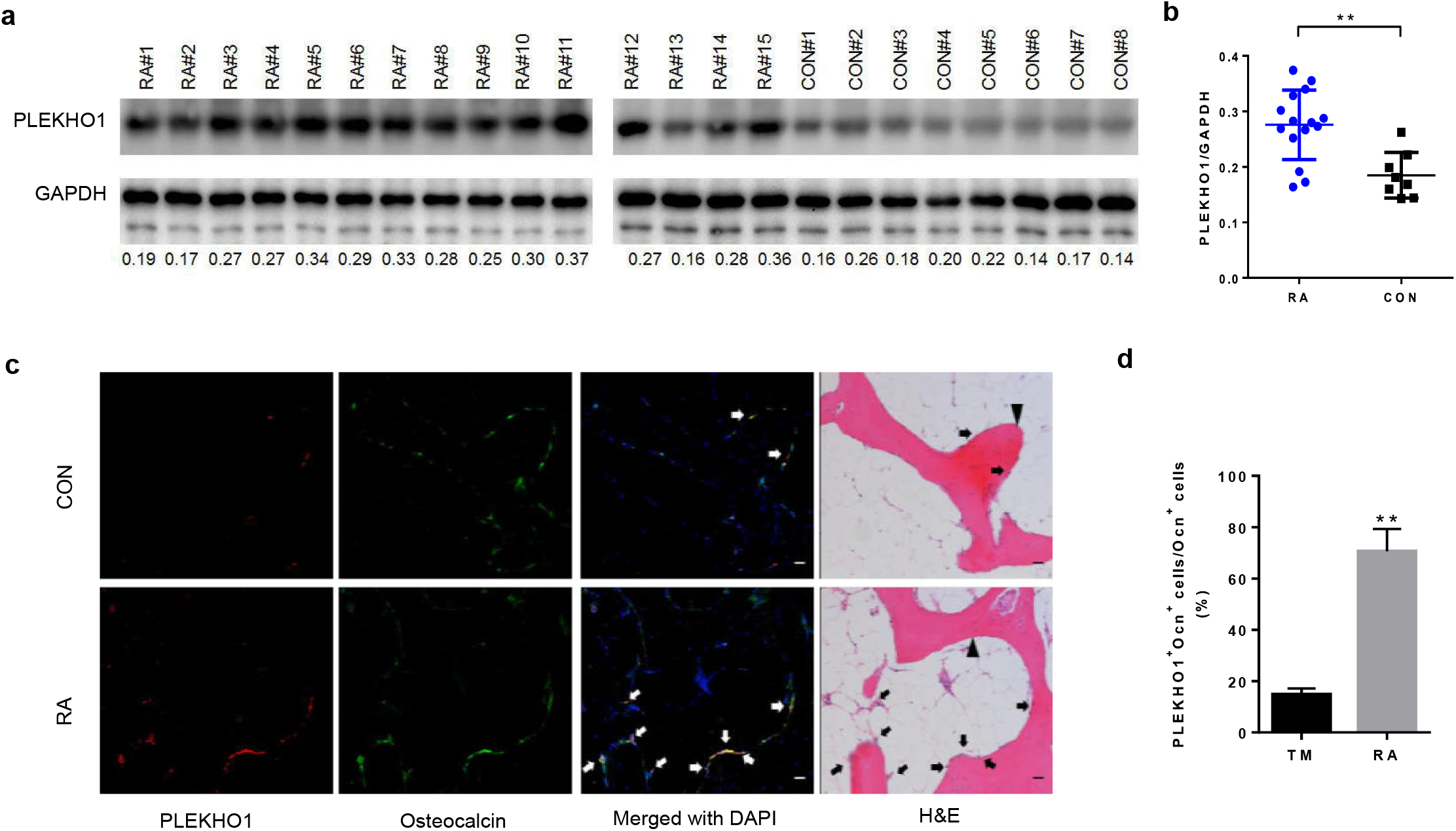
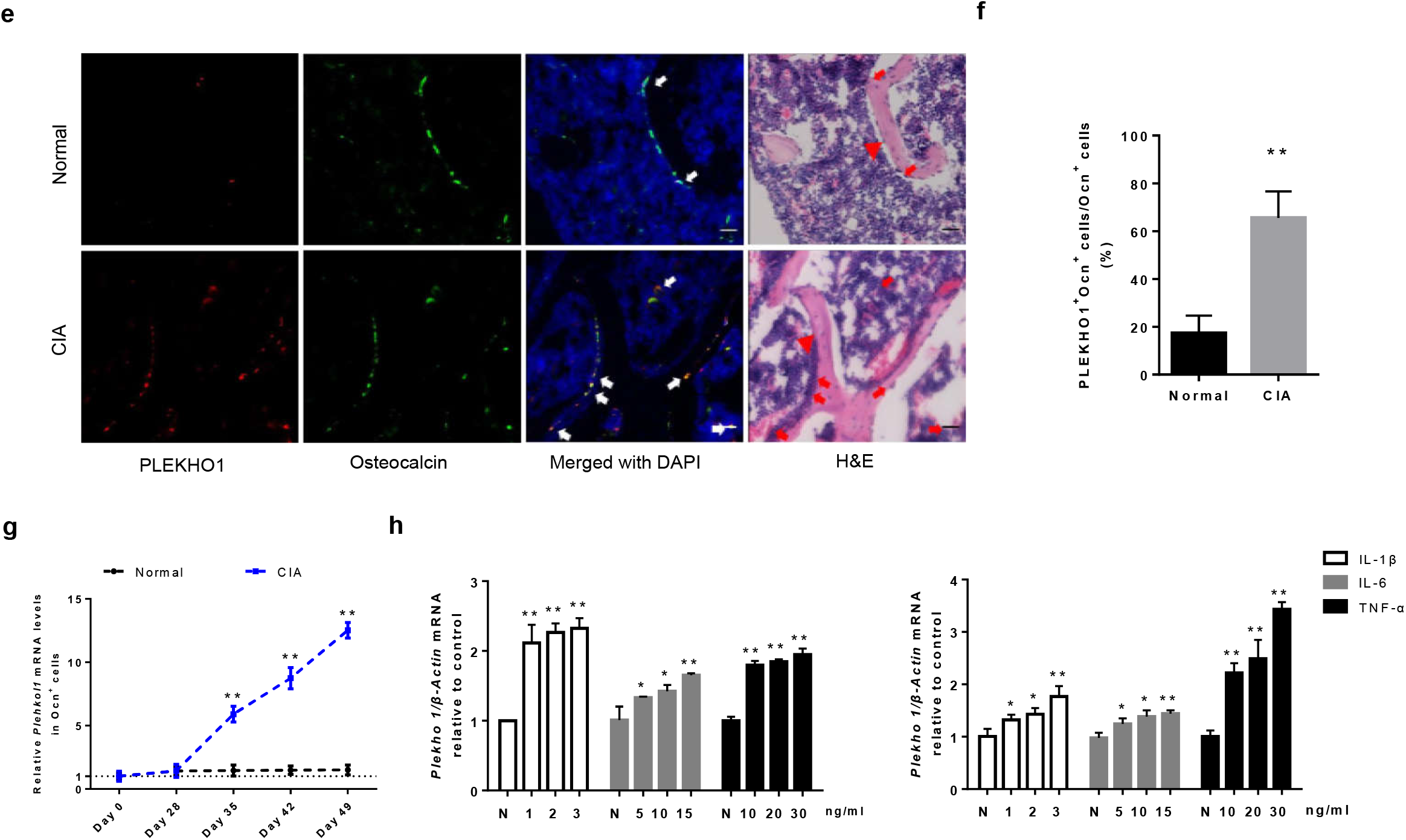
Highly expressed PLEKHO1 in osteoblasts of RA patients and CIA mice. (**a,b**) Comparison of PLEKHO1 levels in bone tissues from knee joint between RA and TM patients underwent knee joint replacement surgery by western blot. **a**, Representative electrophoretic bands of the samples from fifteen RA patients and eight TM patients are shown. The numbers below the bands represent a semiquantitative value. **b**, The semiquantitative data of the protein levels. (**c,d**) Comparison of PLEKHO1 expression (red) within osteocalcin positive (Ocn^+^) osteoblasts (green) in bone tissues from knee joint between RA and TM patients by immunofluorescence analysis. **c**, Representative fluorescent micrographs of the samples from five RA patients and five TM patients are shown. Merged images with DAPI staining showed co-staining of PLEKHO1 and Ocn^+^ osteoblasts (arrows, yellow). H&E staining of the same sections is shown, and black arrowheads point to bone-formation surfaces, enriched by those cells with pink, which is a merged color of green (osteoblast marker) and blue (DAPI staining for nuclei) in the immunofluorescence staining. Scale bars, 20 μm. **d**, Comparison of the percentage of cells co-expressing Ocn and PLEKHO1 in Ocn^+^ cells from bone tissues of knee joint between RA patients and TM patients. (**e,f**) Comparison of PLEKHO1 expression (red) within Ocn^+^ osteoblasts (green) in bone tissues of ankle joint from hind paws of CIA and normal mice on day 49 after primary immunization by immunofluorescence analysis. **e**, Representative fluorescent micrographs of the bone samples from ten CIA mice and ten normal mice are shown. Merged images with DAPI staining showed co-staining of PLEKHO1 and Ocn^+^ osteoblasts (arrows, yellow). H&E staining of the same sections is shown, and red arrowheads point to bone-formation surfaces. Scale bars: 20 μm. **f**, Comparison of the percentage of cells co-expressing Ocn and PLEKHO1 in Ocn^+^ cells from bone tissues of ankle joint from hind paws between CIA mice and normal mice on day 49 after primary immunization. (**g**) Time course changes of *Plekho1* mRNA level in Ocn^+^ osteoblasts from CIA and normal mice by laser-capture microdissection (LCM) combined with qPCR. (**h**) Levels of *Plekho1* mRNA induced by proinflammatory cytokines (IL-1β, IL-6 and TNF-α) in human osteoblast–like cell line MG-63 (left) and mouse osteoblast–like cell line MC3T3E1 (right) by real time PCR analysis. N: no cytokines, served as control. Note: RA, rheumatoid arthritis; TM, severe trauma. All data are the mean ± s.d. *P < 0.05, **P < 0.01. Two-way ANOVA with Bonferroni posttests was performed. Comparisons between two groups were performed using a Student’s t test.

To further investigate the dynamic changes of osteoblastic PLEKHO1 in RA progression, we observed the time course changes in PLEKHO1 expression within osteoblasts in CIA mice, a classic animal model of RA. In line with the above findings in RA patients’ bone specimens, we also found the percentage of cells co-expressing PLEKHO1 and Ocn in Ocn+ cells was higher in bone tissues of ankle joint in the hind paws from CIA mice on day 49 after primary immunization compared to those normal controls (**Fig. 1e, 1f**). Then, we collected Ocn+ cells of ankle joint from CIA group by laser capture micro-dissection (LCM) at different time point after immunization. Further real time PCR analysis indicated that the *Plekho1* mRNA level in Ocn+ cells of ankle joint from CIA group was increased over time from day 28 and significantly higher than that in control group on day 35, 42 and 49, respectively (**Fig. 1g**).

These data suggested that local inflammatory environment might lead to the aberrant upregulation of PLEKHO1 in osteoblasts of joint bone tissues. To test this hypothesis *in vitro,* we stimulated human osteoblast−like cell line MG-63 and mouse osteoblast–like cell line MC3T3E1 with the recombinant proinflammatory cytokines IL-1β, IL-6 and TNF-α, and found higher levels of *Plekho1mRNA* in these cells (**Fig. 1h**).

### Osteoblastic Plekho1 deletion attenuates joint inflammation in inflammatory arthritis mice

To test whether deletion of osteoblastic Plekho1 would attenuate joint inflammation during inflammatory arthritis, we generated the heterozygous mice carrying the mutant allele with LoxP sites harboring exon 3 to exon 6 of *Plekho1* gene *(Plekho1*^*fl*/-^). These mice were then crossed with *Osx-Cre* mice to generate the osteoblast-specific *Plekho1* conditional knockout *(Plekho1*_osx_^-/-^) mice (Supplementary Fig. 1a-b). Real time PCR analysis showed that the Plekho1 mRNA level was hardly detected in Ocn^+^ cells (osteoblasts) from *Plekho1*_osx_^-/-^ mice when compared to control littermates (hereafter WT mice, including *Plekho1*^*fl/fl*^ mice, *Osx*^+/-^;*Plekho1*^fl/-^ mice, and *Osx*^+/-^ mice), whereas no significant difference in *Plekho1* mRNA level within Ocn^-^ cells (non-osteoblasts) was found between *Plekho1*_osx_^-/-^ mice and WT mice (**Supplementary Fig. 1c**).

Subsequently, we examined the joint inflammation changes of *Plekho1*_osx_^-/-^ mice after induction with type II chicken collagen. We found that *Plekho1*_osx_^-/-^-CIA mice showed obvious attenuation in joint inflammation. The arthritis score in *Plekho1*_osx_^-/-^-CIA mice was significantly lower than that in WT mice from day 37 to day 49 after first immunization (**Fig. 2a**). Histological evaluation showed that the score for inflammation in ankle joint from the hind paws was significantly lower in *Plekho1*_osx_^-/-^-CIA mice than that in WT-CIA mice on day 49 after primary immunization (**Fig. 2b, 2c**). Moreover, IL-1β and IL-6 levels in ankle joint of hind paws from *Plekho1*_osx_^-/-^-CIA mice were also remarkably lower on day 49 when compared with the WT-CIA mice (**Fig. 2d**). As previous studies regarded that joint inflammation progression was mainly caused by the activated fibroblast-like synoviocytes (FLS) and immune cells including infiltrating lymphocytes and neutrophils[12], we then detected the cross-talk between osteoblasts and FLS, CD4+T lymphocytes as well as neutrophils by using an *in vitro* co-culture system which could experimentally simulate the temporo-spatial complexity of the *in vivo* cross-talk between osteoblasts and inflammatory cells in the joint. The osteoblasts were differentiated from the primary osteoblast precursor cells isolated from the calvarial bone of newborn *Plekho1*_osx_^-/-^ mice and WT littermates in osteogenic medium and confirmed to express osteocalcin. The FLS, CD4+T lymphocytes and neutrophils were obtained from WT mice by collagenase digestion and specific magnetic beads-based selection, respectively. Thereafter, the FLS, CD4+T lymphocytes or neutrophils were co-cultured with *Plekho1*_osx_^-/-^ or WT osteoblasts, respectively. After co-culture, the cells were collected for real time PCR analysis. The results showed that under TNF-α stimulation, the IL-1β and IL-6 mRNA levels in the CD4+T lymphocytes and FLS were remarkably decreased after co-culture with osteoblasts from *Plekho1*_osx_^-/-^ mice when compared with the control group (**Fig. 2e, 2f**). The migration of neutrophils was also inhibited after co-culture with osteoblasts from *Plekho1*_osx_^-/-^ mice (**Fig. 2g**). These novel results pointed to an important role of osteoblastic PLEKHO1 in modulating local inflammatory response in RA.

**Figure 2.**
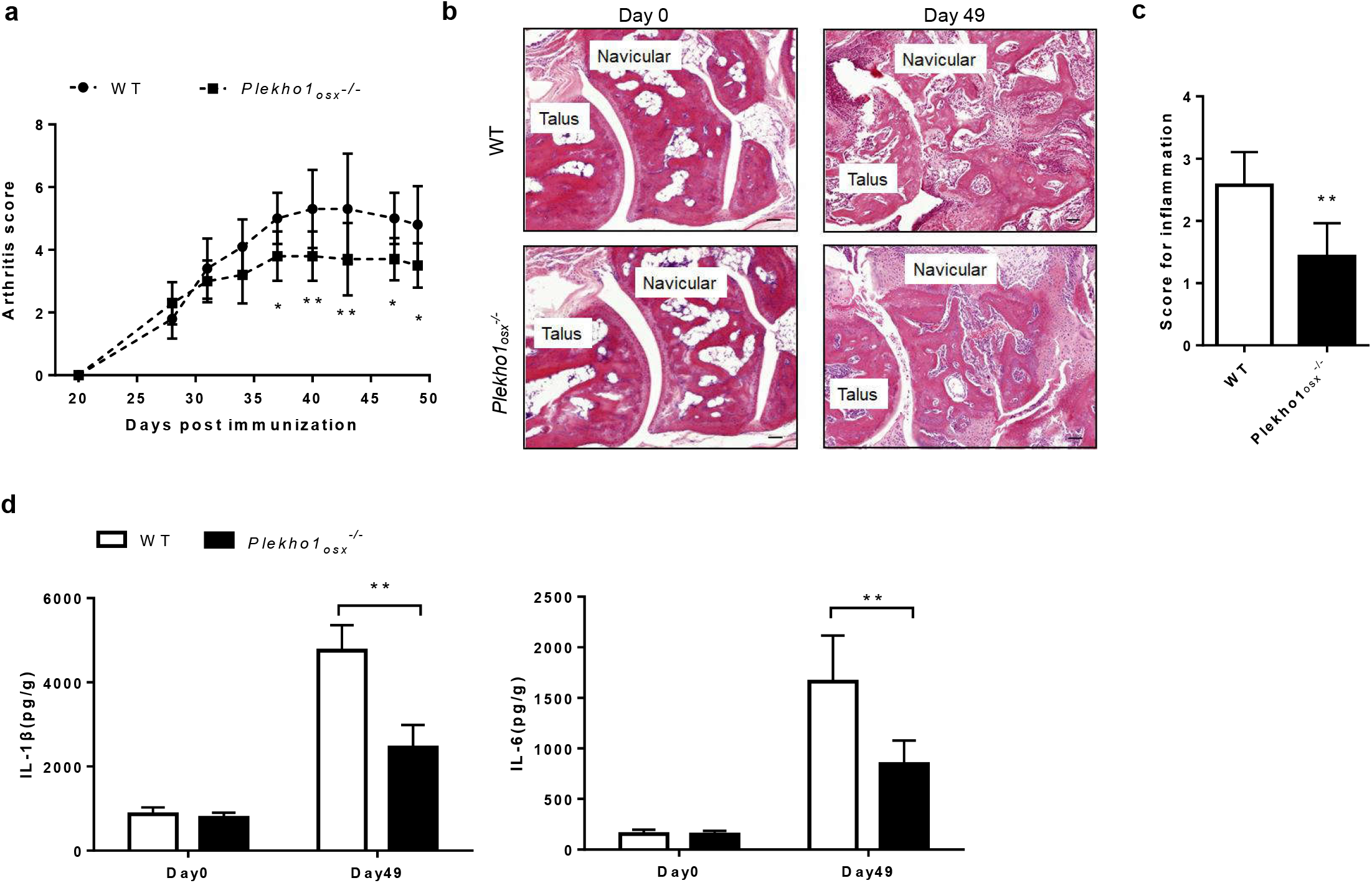
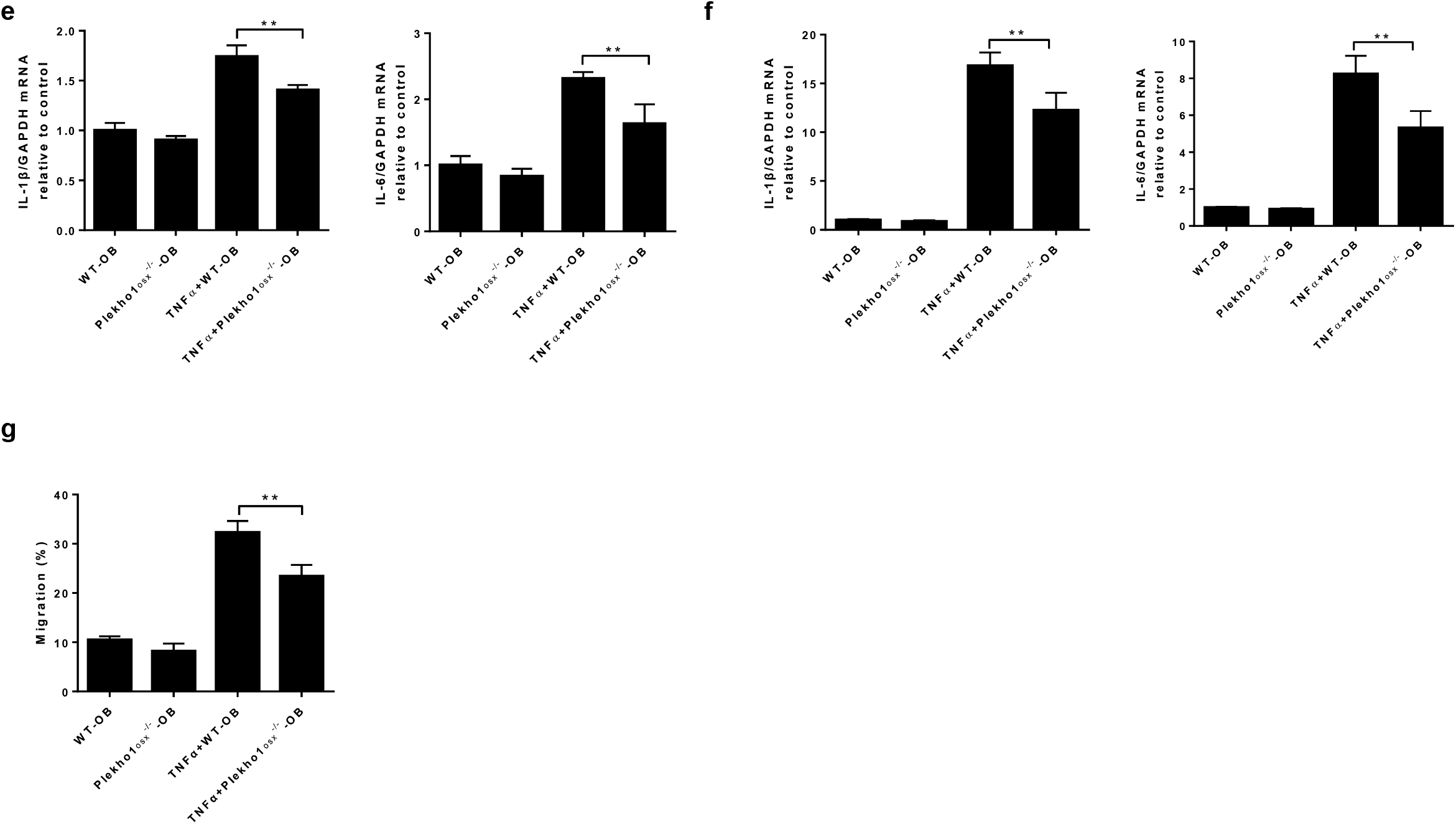
Genetic deletion of osteoblastic Plekho1 leads to amelioration of joint inflammation in CIA mice. (**a**) Time course changes in arthritis score from *Plekho1_osx_*^-/-^ mice and WT mice (including *Plekho1^fl/fl^* mice, *Osx^+/-^;Plekho1^fl/-^* mice, and *Osx*^+/-^ mice) after induction with type II chicken collagen. (**b**) Representative histological images in the ankle joint of hind paws from *Plekho1_osx_*^-/-^-CIA mice and WT-CIA mice on day 49 after primary immunization. Scale bars, 50μm. (c) Histological inflammation score in the ankle joint of hind paws from *Plekho1_osx_*^-/-^-CIA mice and WT-CIA mice on day 49 after primary immunization. (**d**) Time course changes of IL-1 β and IL-6 levels in the ankle joint of hind paws from *Plekho1_osx_*^-/-^-CIA mice and WT-CIA mice by ELISA examination. (**e**) IL-1 β and IL-6 mRNA levels in CD4+T lymphocytes after co-culture with osteoblasts from WT mice and *Plekho1_osx_*^-/-^ mice. (**f**) IL-1 β and IL-6 mRNA levels in fibroblast-like synoviocytes after co-culture with osteoblasts from WT mice and *Plekho1_osx_*^-/-^ mice. (**g**) Migration of neutrophils after co-culture with osteoblasts from WT mice and *Plekho1_osx_*^-/-^ mice. All data are the mean ± s.d. *n* = 10 per group. *P < 0.05, **P < 0.01. For **a**, Two-way ANOVA with subsequent Bonferroni posttests was performed. For **c & d**, a Student’s t test was performed. For **e-g**, a one-way ANOVA with subsequent Tukey’s multiple comparisons test was performed.

As our previous studies demonstrated that osteoblastic PLEKHO1 negatively regulated bone formation during aging-related bone loss and osteoporosis[8, 9], we consistently examined the bone phenotypes of *Plekho1*_osx_^-/-^ -CIA mice. MicroCT analysis showed a better organized architecture and a higher bone mass in *Plekho1*_osx_^-/-^-CIA mice compared to WT-CIA mice on day 49 after primary immunization, both in cortical bone and trabecular bone (**Supplementary Fig. 2a**). The values of bone mineral density (BMD), trabecular fraction (BV/TV), trabecular thickness (Tb.Th) and trabecular number (Tb.N) of the tibial bones were gradually decreased from baseline (day 0) in WT-CIA mice and *Plekho1*_osx_^-/-^-CIA mice. However, *Plekho1*_osx_^-/-^-CIA mice showed a slow decrease in these parameters on day 49 compared to the rapid decrease in the control mice (**Supplementary Fig. 2b**). Dynamic bone histomorphometric analysis showed a larger width between the xylenol and the calcein labeling bands in *Plekho1*_osx_^-/-^-CIA mice compared to WT-CIA mice (**Supplementary Fig. 2c**).The values of mineral apposition rate (MAR), bone formation rate (BFR/BS), osteoblasts surface (Ob.S/BS) and osteoblast number (Ob.N/B.Pm) were also decreased slower in *Plekho1*_osx_^-/-^-CIA mice than those in WT-CIA mice on day 49 after primary immunization. The values of osteoclasts surface (Oc.S/BS) and osteoclast number (Oc.N/B.Pm) showed a slower increase in *Plekho1*_osx_^-/-^-CIA mice than those in WT-CIA mice on day 49 after primary immunization (**Supplementary Fig. 2d**). These results demonstrated that osteoblastic PLEKHO1 indeed negatively regulated bone formation in CIA mice as well.

In addition, to further evaluate the role of osteoblastic PLEKHO1 in local inflammation regulation during RA, we induced *Plekho1*_osx_^-/-^ mice with K/BxN serum, which developed an acute arthritis and was more sensitive in mice with C57BL/6 background. Of note, we again observed less joint inflammation in *Plekho1*_osx_^-/-^-STA mice, which was reflected by lower arthritis score from day 8 and lower histological inflammation score in ankle joint of hind paws at day 12 in *Plekho1*_osx_^-/-^-STA mice compared to the control mice (**Supplementary Fig. 3a-c**). Moreover, IL-1β and IL-6 levels in ankle joint of hind paws from *Plekho1*_osx_^-/-^-STA mice were also remarkably lower on day 12 when compared with the WT-STA mice (**Supplementary Fig. 3d**).

### Overexpressing Plekho1 only in osteoblasts exacerbates joint inflammation in inflammatory arthritis mice

To exclude the interference of PLEKHO1 in other cells, we created another mouse strain carrying the ROSA26-PCAG-STOP^fl^-*Plekho1*-eGFP allele, and crossed them with the Osx-Cre mice to generate the osteoblast-specific Plekho1 overexpressing (*Plekho1*_osx_ *Tg*) mice that overexpressing PLEKHO1 in osteoblasts. Then, we crossed *Plekho1*^-/-^ mice with *Plekho1*_osx_ *Tg* mice to obtain mice that express high Plekho1 exclusively in osteoblasts *(Plekho1*^-/-^/*Plekho1*_osx_ *Tg* mice) (**Supplementary Figure 4a, 4b**). Real time PCR analysis showed that *Plekho1*^-/-^/*Plekho1*_osx_ *Tg* mice had remarkably higher *Plekho1* mRNA levels in Ocn^+^ cells (osteoblasts), at the same time, had little *Plekho1* mRNA levels in Ocn^-^ cells (non-osteoblasts) when compared to control littermates (hereafter WT mice, including *Osx*^+/-^;*Plekho1*^*fl*/-^ mice and *Osx*^+/-^;*Plekho1*^*fl/fl*^ mice). *Plekho1*^-/-^ mice had hardly detected Plekho1 mRNA levels in both Ocn^+^ cells and Ocn^-^ cells (**Supplementary Figure 4c**). Then, we induced these mice with CIA. We found that exclusively high PLEKHO1 expression in osteoblasts exacerbated the disease severity and joint inflammation progression, which were shown by increased arthritis score from day 35 after primary immunization and higher pathological inflammation score of ankle joint form the hind paws in *Plekho1*^-/-^/*Plekho1_osx_ Tg*-CIA mice compared to WT-CIA mice on day 49 after primary immunization (**Fig. 3a-c**). IL-1β and IL-6 levels in ankle joint of hind paws from *Plekho1*^-/-^/*Plekho1_osx_* Tg-CIA mice were also obviously higher when compared to WT-CIA mice on day 49 after primary immunization (**Fig. 3d**). In addition, we isolated primary osteoblast precursor cells from calvarial bone of newborn *Plekho1_osx_ Tg* mice and WT littermates, these precursor cells were induced maturation for co-culture with CD4+T lymphocytes, FLS and neutrophils isolated from WT mice, respectively. we found that under TNF-α stimulation, the IL-1β and IL-6 mRNA levels in CD4+T lymphocytes and FLS were higher after co-culture with osteoblasts from *Plekho1_osx_ Tg* mice when compared with the control group (**Fig. 3e, 3f**). The migration of neutrophils was also enhanced after co-culture with osteoblasts from *Plekho1_osx_ Tg* mice (**Fig. 3g**).

**Figure 3.**
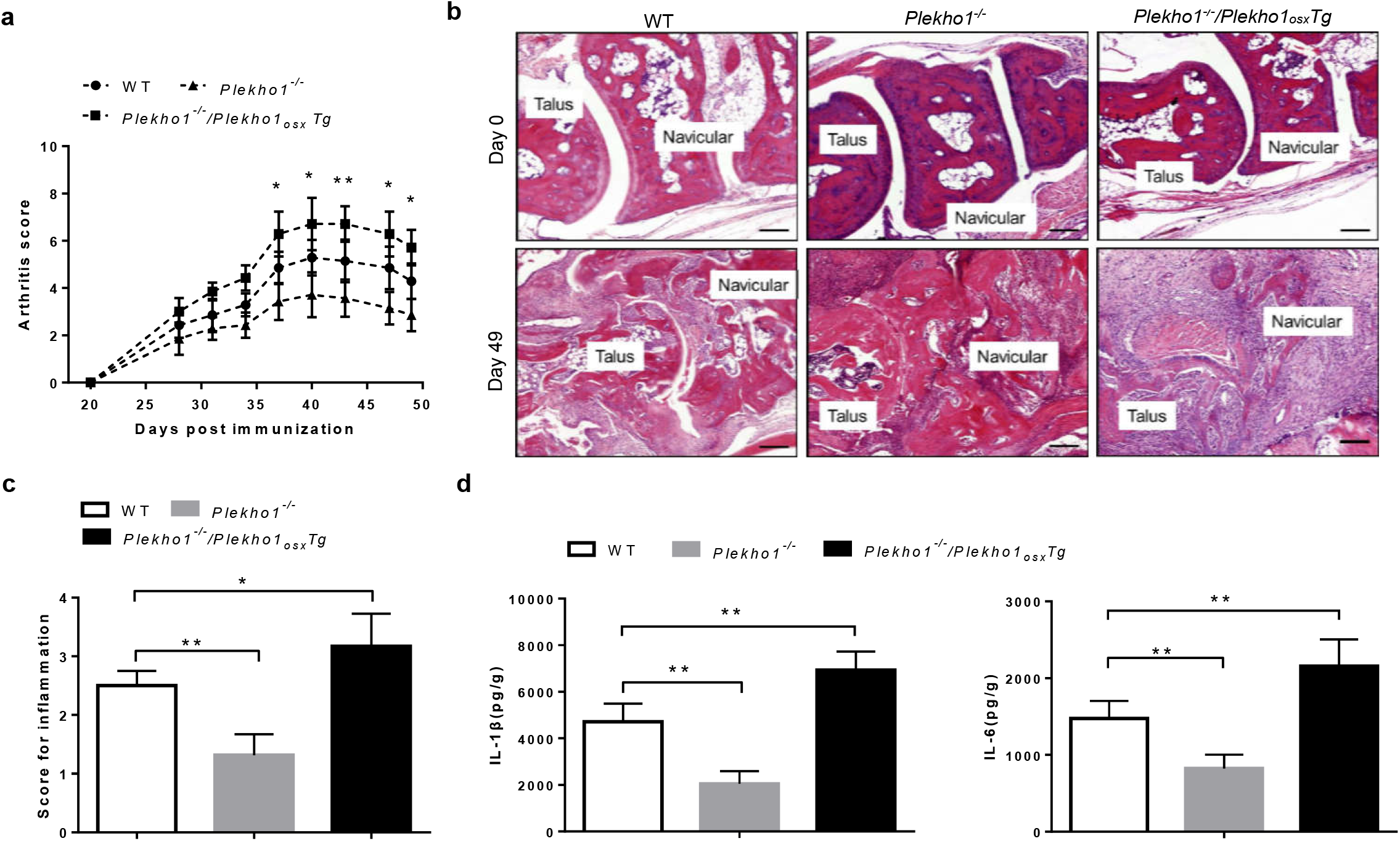
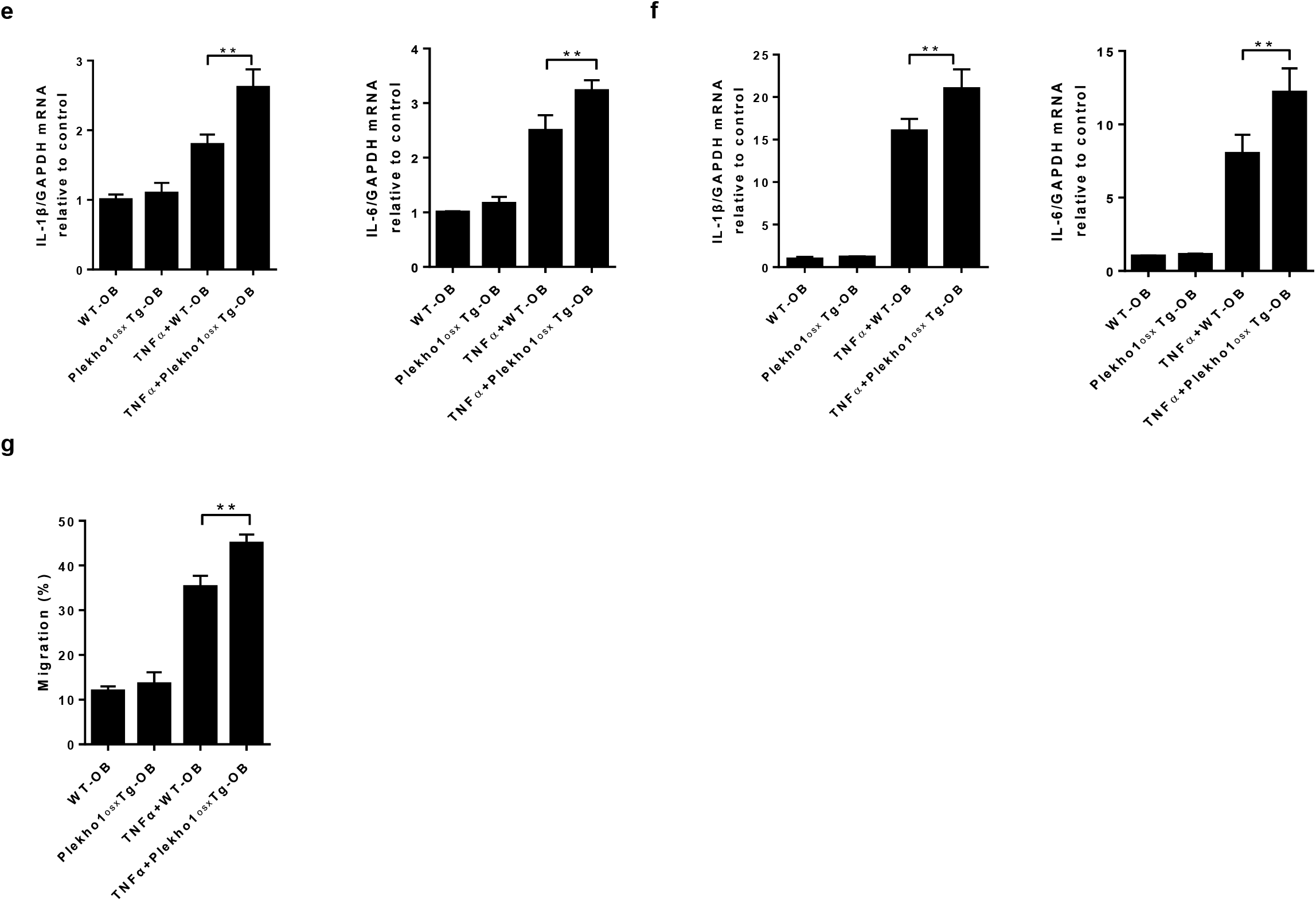
Overexpressed Plekho1 in osteoblasts exacerbates inflammation in *Plekho1*^-/-^/*Plekho1_osx_ Tg* mice with CIA. (**a**) Disease progression assessed by arthritic score in WT-CIA, *Plekho1*^-/-^-CIA and *Plekho1*^-/-^/*Plekho1_osx_* Tg-CIA mice. *P < 0.05, **P < 0.01, compared to WT-CIA group. (**b**) Representative histological images in the ankle joint of hind paws from WT-CIA, *Plekhot1*^-/-^-CIA and *Plekho1*^-/-^/*Plekho1_osx_ Tg*-CIA mice. Scale bars, 50μm. (**c**) Histological inflammation score in the ankle joint of hind paws from WT-CIA, *Plekho1*^-/-^-CIA and *Plekho1*^-/-^/*Plekho1_osx_ Tg*-CIA mice. (**d**) IL-1β and IL-6 levels in the ankle joint of hind paws from WT-CIA, *Plekho1*^-/-^-CIA and *Plekho1*^-/-^/*Plekho1_osx_ Tg*-CIA mice on day 49 after primary immunization by ELISA examination. (**e**) IL-1 β and IL-6 mRNA levels in CD4+T lymphocytes after co-culture with osteoblasts from WT mice and *Plekho1_osx_ Tg* mice. (**f**) IL-1 β and IL-6 mRNA levels in fibroblast-like synoviocytes after co-culture with osteoblasts from WT mice and *Plekho1_osx_ Tg* mice. (**g**) Migration of neutrophils after co-culture with osteoblasts from WT mice and *Plekho1_osx_ Tg* mice. All data are the mean ± s.d. *n* = 9 per group. *P < 0.05, **P < 0.01. For **a**, a two-way ANOVA with subsequent Bonferroni posttests was performed. For **c-g**, a one-way ANOVA with subsequent Tukey’s multiple comparisons test was performed.

We also induced *Plekho1^-/-^/Plekho1_osx_ Tg* mice with K/BxN serum. Consistently, we again observed more joint inflammation in *Plekho1^-/-^/Plekho1_osx_ Tg*-STA mice, which was reflected by quicker increased arthritis score and higher histological inflammation score in the ankle joint of hind paws in *Plekho1^-/-^/Plekho1_osx_ Tg*-STA mice compared to the control littermates (**Supplementary Fig. 5a-c**). IL-1β and IL-6 levels in ankle joint of hind paws from *Plekho1^-/-^/Plekho1_osx_ Tg*-STA mice were significantly higher on day 12 when compared with the WT-STA mice (**Supplementary Fig. 5d**). These results further demonstrated the important role of osteoblastic PLEKHO1 in modulating joint inflammation in RA.

### PLEKHO1 is required for TRAF2-mediated RIP1 ubiquitination to activate NF-kB for inducing inflammatory cytokines production in osteoblasts

Then, to detect whether PLEKHO1 could affect TNF-α-stimulated osteoblasts releasing inflammatory effectors so as to modulate inflammation, we used a mouse osteoblast–like cell line MC3T3E1. We found that after Plekho1 silencing by Plekho1 siRNA, the IL-1β and IL-6 production in supernatant of TNF-α-stimulated MC3T3E1 cells were remarkably lower than the random siRNA group (**Fig. 4a**). To further confirm it *in vivo,* we crossed hTNFtg mice, which could spontaneously develop arthritis at the age of four week due to the overexpressed human TNF transgene, with *Plekho1_osx_*^-/-^ mice to obtain *Plekho1_osx_*^-/-^/hTNFtg mice. Notably, *Plekho1_osx_*^-/-^/hTNFtg mice showed a distinct alleviation of disease severity. The arthritis score in *Plekho1_osx_*^-/-^/hTNFtg mice was significantly lower than that in hTNFtg mice from 7 week to 10 week after birth (**Supplementary Figure 6a**). Histological analysis also indicated that joint inflammation was significantly ameliorated in *Plekho1_osx_*^-/-^/hTNFtg mice compared to hTNFtg mice, which were reflected by lower histological inflammation score at week 10 (**Supplementary Figure 6b, 6c**). These results implied that osteoblastic PLEKHO1 on joint inflammation regulation might be related with TNF-α involved signal mechanism. As previous studies have demonstrated that NF-κB signal pathway played an important role in TNF-α induced inflammatory cytokines production[13], we therefore detected the change of related proteins in this signal pathway after specifically silencing Plekho1 by Plekho1 siRNA. We found that IKKα/β and p65 phosphorylation as well as RIP1 ubiquitination were inhibited under TNF-α stimulation in Plekho1-silenced MC3T3E1 cells, although no significantly change on production of IKKβ and p65 (**Fig. 4b-d**).

**Figure 4.**
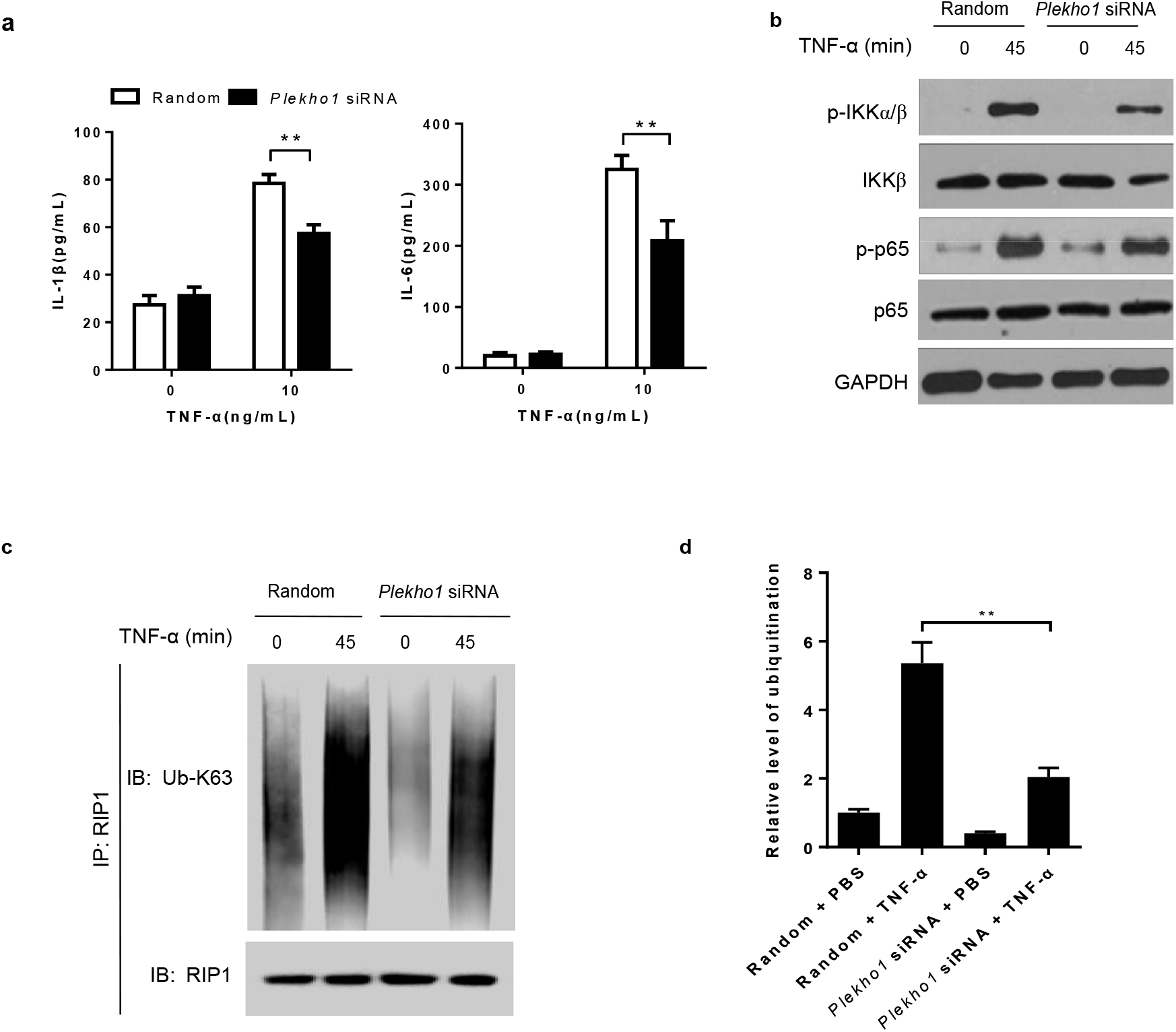
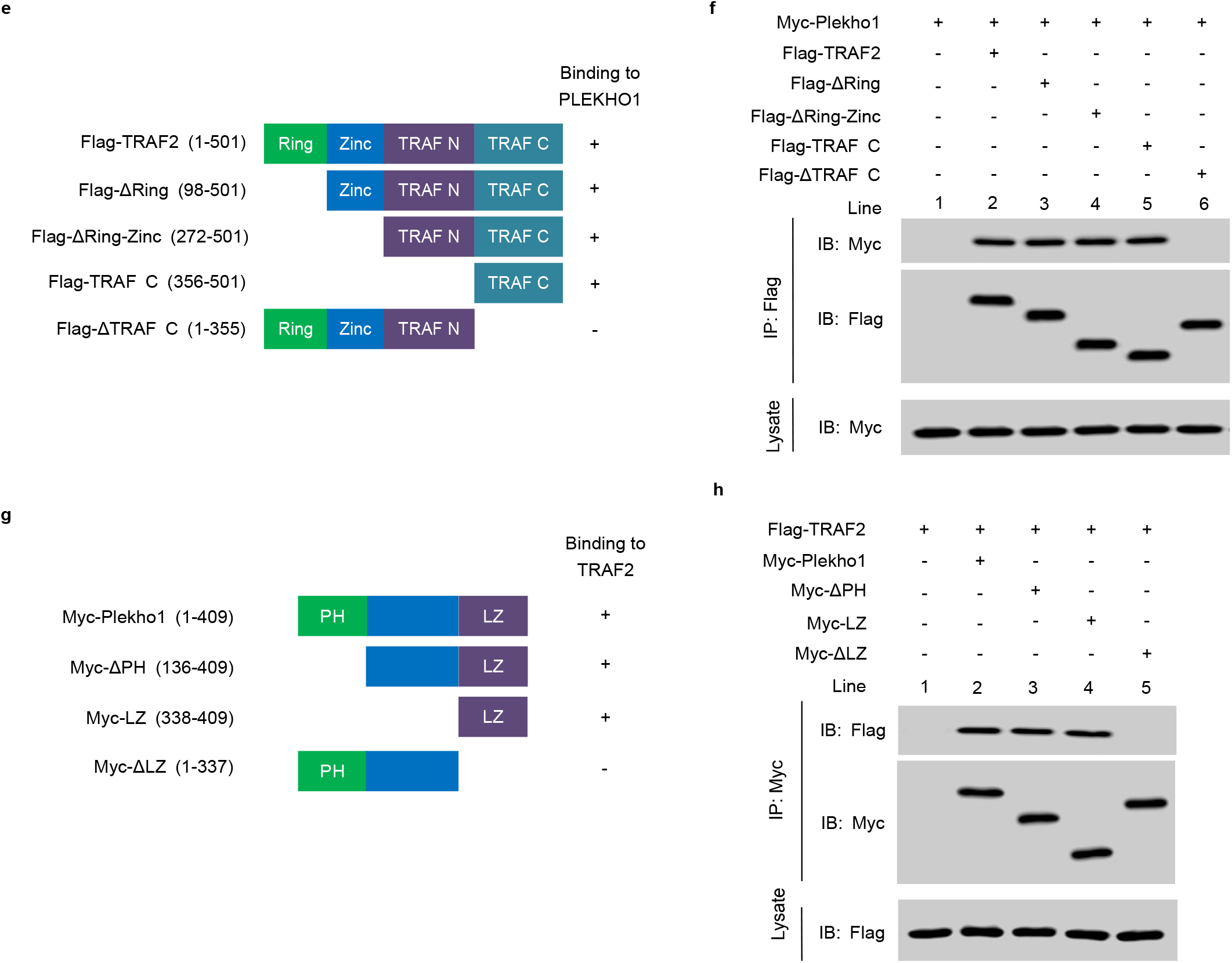
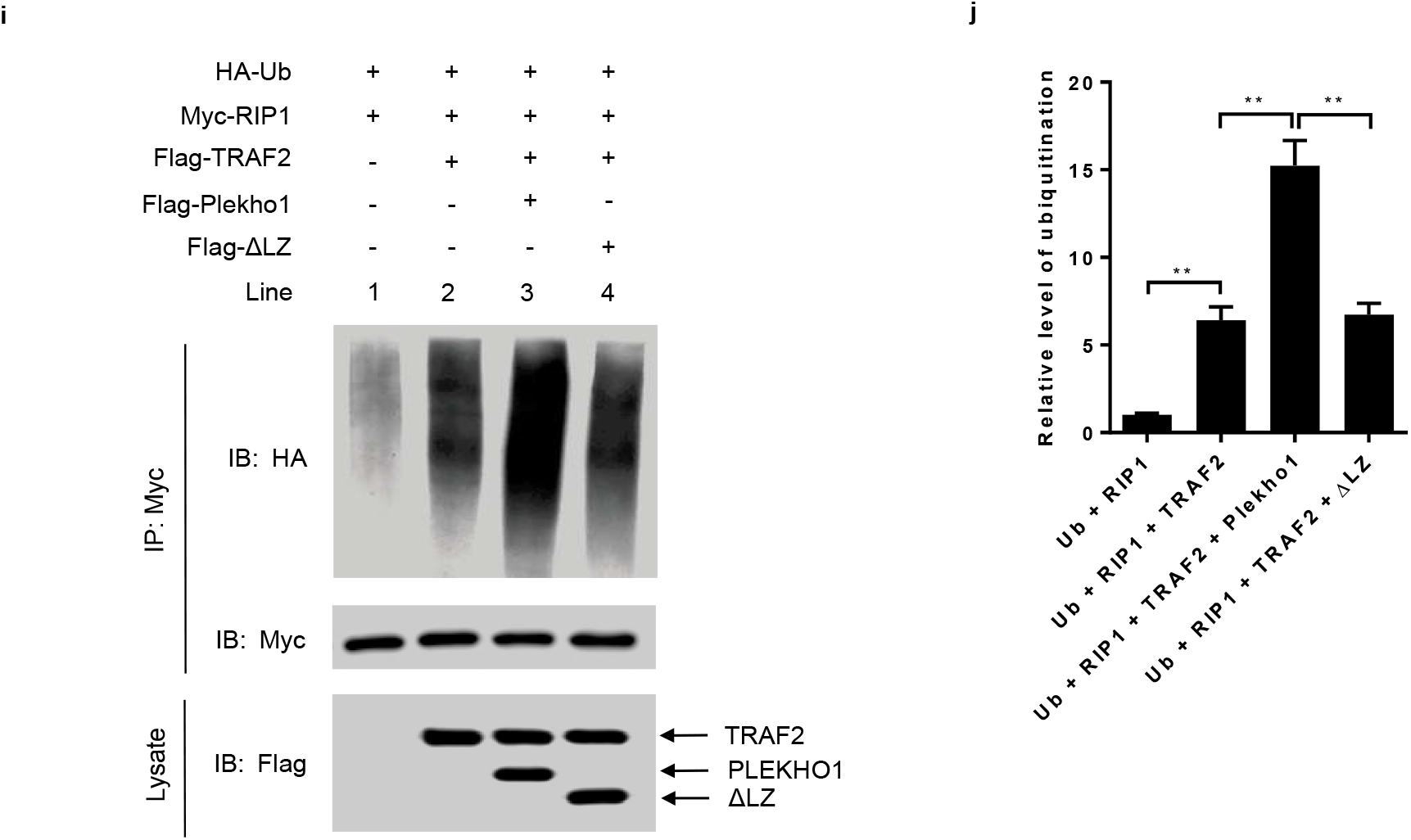
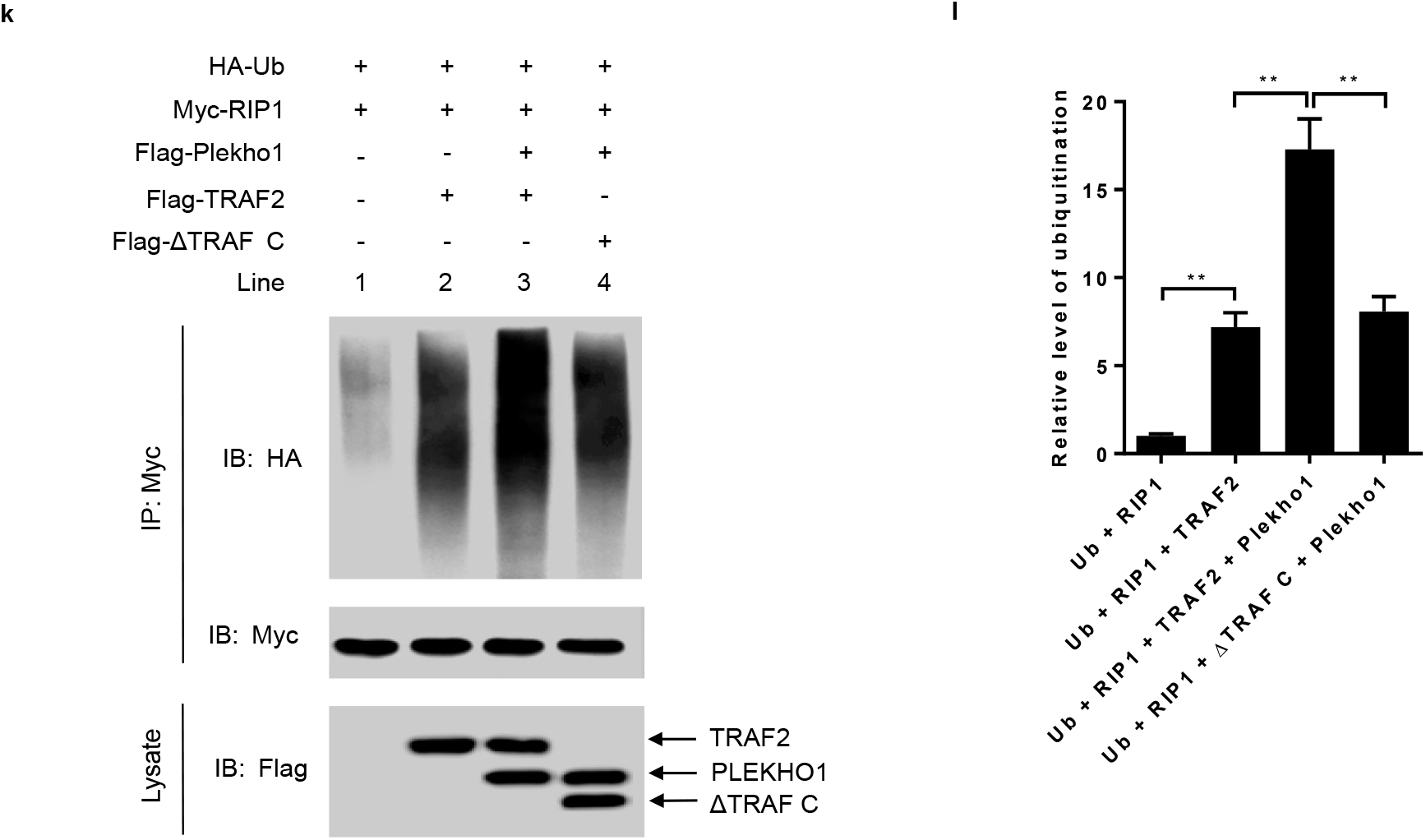
PLEKHO1 interacts with TRAF2 to promote the TRAF2-mediated ubiquitination of RIP1. (**a**) Levels of inflammatory cytokines (IL-1 β, IL-6) produced by MC3T3E1 cells transfected with *Plekho1* siRNA or random siRNA and subjected to TNF-α (10 ng/ml). (**b**) MC3T3E1 cells transfected with *Plekho1* siRNA or random siRNA were stimulated with TNF-α (10 ng/ml) for 45min, and harvested for detection of IKKα/β and p65 phosphorylation. (**c**) MC3T3E1 cells transfected with *Plekho1* siRNA or random siRNA were stimulated with TNF-α (10 ng/ml) for 45min, and harvested for detection of RIP1 ubiquitination. (**d**) Quantification of relative level of RIP1 ubiquitination. (**e, f**) Immunoprecipitation using anti-Flag antibody to examine the PLEKHO1-TRAF2 interaction and mapping of the PLEKHO1-interacting domain in TRAF2. **e**, The Flag-tagged full-length TRAF2 (Flag-TRAF2) and various truncated mutants (Flag-ΔRing, Flag-ΔRing-Zinc, Flag-TRAF N and Flag-ΔTRAF C) were shown with amino acid numbers. **f**, HEK293T cells were transfected with Myc-Plekho1 and Flag-TRAF2 or truncated mutants as indicated. Full-length TRAF2 or truncated mutants was immunoprecipitated by anti-Flag antibody. PLEKHO1 in immunoprecipitates or whole-cell lysate was detected by anti-Myc antibody. (**g, h**) Immunoprecipitation using anti-Myc antibody to examine the PLEKHO1-TRAF2 interaction and mapping of the TRAF2-interacting domain in PLEKHO1. **g**, The Myc-tagged full-length Plekho1 (Myc-Plekho1) and various truncated mutants (Myc-ΔPH, Myc-LZ, and Myc-ΔLZ) were shown with amino acid numbers. **h**, HEK293T cells were transfected with Flag-TRAF2 and Myc-Plekho1 or truncated mutants as indicated. Full-length PLEKHO1 or truncated mutants was immunoprecipitated by anti-Myc antibody. TRAF2 in immunoprecipitates or whole-cell lysate was detected by anti-Flag antibody. (**i**) *In vivo* ubiquitination assay of RIP1 in HEK293T cells transfected with Myc-RIP1, HA-Ubiquitin (HA-Ub) and Flag-TRAF2, along with Flag-Plekho1 or Flag-ΔLZ. Ubiquitination of RIP1 was detected in Myc immunoprecipitates. (**j**) Quantification of relative level of RIP1 ubiquitination. (**k**) *In vivo* ubiquitination assay of RIP1 in HEK293T cells transfected with Myc-RIP1, HA-Ub and Flag-Plekho1, along with Flag-TRAF2 or Flag-ΔTRAF C. Ubiquitination of RIP1 was detected in Myc immunoprecipitates. (**l**) Quantification of relative level of RIP1 ubiquitination. Note: IP, Immunoprecipitation; IB, immunoblotting. Data are representative of three independent experiments. **P<0.01. Comparisons between two groups were performed using a Student’s t test.

Previous studies demonstrated that TNF-α up-regulated the expression of inflammatory mediators through TRAF2-mediated NF-κB activation in MC3T3E1 cells[14]. In addition, PLEKHO1 could interact with the member of TRAF family to regulate the downstream signal[6], we therefore hypothesized that PLEKHO1 might interact with TRAF2 to promote the activation of NF-kB signal pathways. TRAF2 contains a ring finger domain, a zinc finger domain, a TRAF N domain and a TRAF C domain[15-17]. To test whether TRAF2 could interact with PLEKHO1 and which domain in TRAF2 was essential for their interaction, we constructed Myc-tagged full-length Plekho1 (Myc-Plekho1), Flag-tagged full-length TRAF2 (Flag-TRAF2) and various TRAF2 truncated mutants including Flag-ΔRing, Flag-ΔRing-Zinc, Flag-TRAF N and Flag-ΔTRAF C (**Fig. 4e**). HEK293T cells were transfected with Myc-Plekho1 and Flag-TRAF2 or truncated mutants. After immunoprecipitation using anti-Flag antibody, we detected that TRAF2 could interact with PLEKHO1 (**Fig. 4f**, Lane 2). However, deletion of TRAF C domain (ΔTRAF C) abolished the interaction between TRAF2 and PLEKHO1 (**Fig. 4f**, Lane 6), whereas other truncated mutants (ΔRing, ΔRing-Zinc and TRAF C) maintained the binding with PLEKHO1 as that of full-length TRAF2 (**Fig. 4f**, Lane 3-5). In addition, PLEKHO1 is composed of an N-terminal pleckstrin homology (PH)-containing domain and a C-terminal leucine zipper (LZ)-containing domain[7, 18]. We also constructed Plekho1 truncated mutants (Myc-ΔPH, Myc-LZ and Myc-ΔLZ) to identify the key domain in PLEKHO1 that interacted with TRAF2 (**Fig. 4g**). Data from immunoprecipitation showed that the Plekho1 mutant with deletion of LZ-containing domain (ΔLZ) could not interact with TRAF2 (**Fig. 4h**, Lane 5), but other Plekho1 mutants could still bound to TRAF2 (**Fig. 4h**, Lane 2-4).

As TRAF2 could ubiquitinate RIP1, which is essential for NF-kB activation[19], we then determined whether PLEKHO1 affected TRAF2-mediated RIP1 ubiquitination. We performed *in vivo* ubiquitination assay in HEK293T cells transfected with Myc-RIP1, HA-Ubiquitin (HA-Ub), Flag-Plekho1 or Flag-ΔLZ and Flag-TRAF2 or Flag-ΔTRAF C. Our results demonstrated that PLEKHO1 promoted the TRAF2-mediated ubiquitination of RIP1 (lane 3 in **Fig. 4i**, lane 3 in **Fig. 4k, Fig. 4j, Fig. 4l**). Nevertheless, PLEKHO1 mutant with deletion of c-terminal LZ-containing domain had no obvious promotive effects on TRAF2-mediated ubiquitination of RIP1 (lane 4 in **Fig. 4i, Fig. 4j**). Disruption of TRAF2-PLEKHO1 interaction by deletion of ΔTRAF C in TRAF2 also diminished the promotive effect of PLEKHO1 on TRAF2-mediated RIP1 ubiquitination (lane 4 in **Fig. 4k, Fig. 4l**). All these results indicated that PLEKHO1 could interact with TRAF2 to promote the TRAF2-mediated ubiquitination of RIP1.

### Osteoblastic PLEKHO1 inhibition alleviates joint inflammation and bone formation reduction in CIA mice

Next, to test whether early inhibition of osteoblastic PLEKHO1 could alleviate joint inflammation and promote bone formation in mice with established arthritis, we used a cross-species *Plekho1* siRNA and a nucleic acid delivery system targeting osteoblasts, which were both established by our group [5, 8]. This cross-species *Plekho1* siRNA could target the four species (human, monkey, rat and mouse). *In vivo* studies demonstrated that periodic intravenous injections of this cross-species *Plekho1* siRNA promoted bone formation and reversed bone loss in the osteoporotic mice. On the other hand, to facilitate delivering this cross-species *Plekho1* siRNA to osteoblasts, we have developed a targeted delivery system – (AspSerSer)_6_-liposome. We confirmed that this targeted delivery system could facilitate *Plekho1* siRNA specifically achieve high gene knockdown efficiency in osteoblasts. Therefore, we then performed six consecutive administration of this cross-species *Plekho1* siRNA encapsulated by the osteoblast-targeting delivery system ((AspSerSer)_6_-liposome-*Plekho1* siRNA, hereafter siRNA) in DBA/1 mice at an interval of one week from day 28 after primary immunization with bovine type II collagen, because arthritis was clinically evident at this time point (**Supplementary Fig. 7a**). We firstly confirmed that the level of *Plekho1* mRNA was significantly downregulated in Ocn^+^ cells of siRNA-treated CIA mice after six-week treatment (on day 70 after primary immunization) (**Supplementary Fig. 7b**). Thereafter, we found that arthritis score was significantly decreased in siRNA-treated CIA mice when compared to mice treated with (AspSerSer)_6_-liposome-non sense siRNA (hereafter NS), (AspSerSer)_6_-liposome (hereafter VEH) or PBS (**Fig. 5a**). Histological evaluation showed that the score of joint inflammation in siRNA-treated CIA mice was also significantly lower when compared to three other control groups after treatment (**Fig. 5b, 5c**). IL-1β and IL-6 levels in ankle joint of hind paws from siRNA-treated CIA mice were also significantly lower when compared with the three other control groups (**Supplementary Fig. 7c**). Subsequently, microCT analysis showed better organized microarchitecture and a higher bone mass in tibial bone of siRNA-treated CIA mice compared to mice treated with NS, VEH or PBS after treatment (**Fig. 5c**). The values of BMD, BV/TV, Tb.N and Tb.Th of the tibial bones had remarkable restoration in siRNA-treated CIA mice, whereas such restoration was not observed in CIA mice treated with NS, VEH or PBS (**Fig. 5d**). Similarly, we found a larger width between the xylenol and calcein labeling bands in the mice treated with siRNA compared to the mice treated with NS, VEH or PBS. The values of MAR, BFR/BS and Ob.S/BS were significantly higher, whereas the value of Oc.S/BS were significantly lower in siRNA group when compared with three other control groups after treatment (**Fig. 5c, Fig. 5e**).

**Figure 5.**
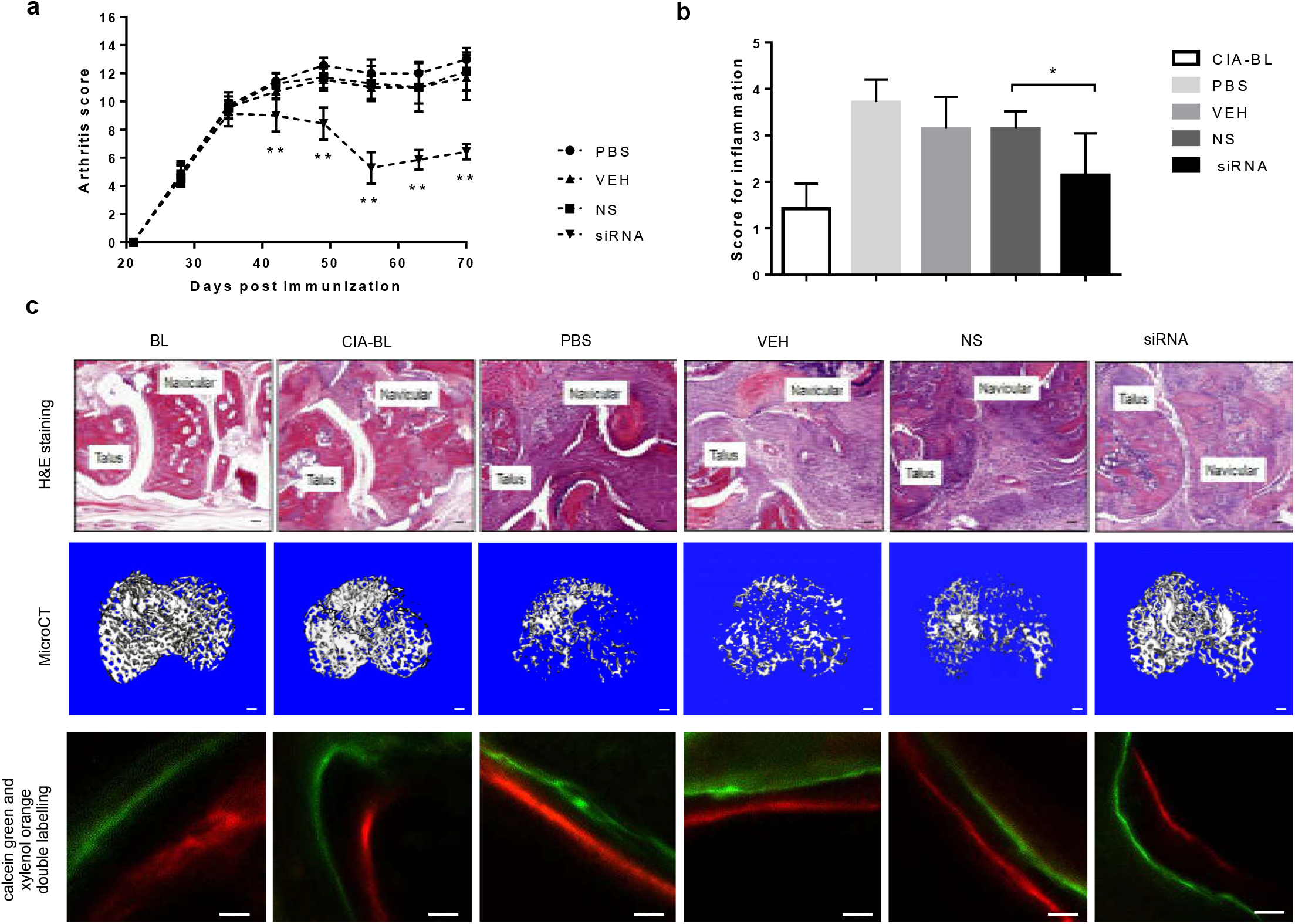
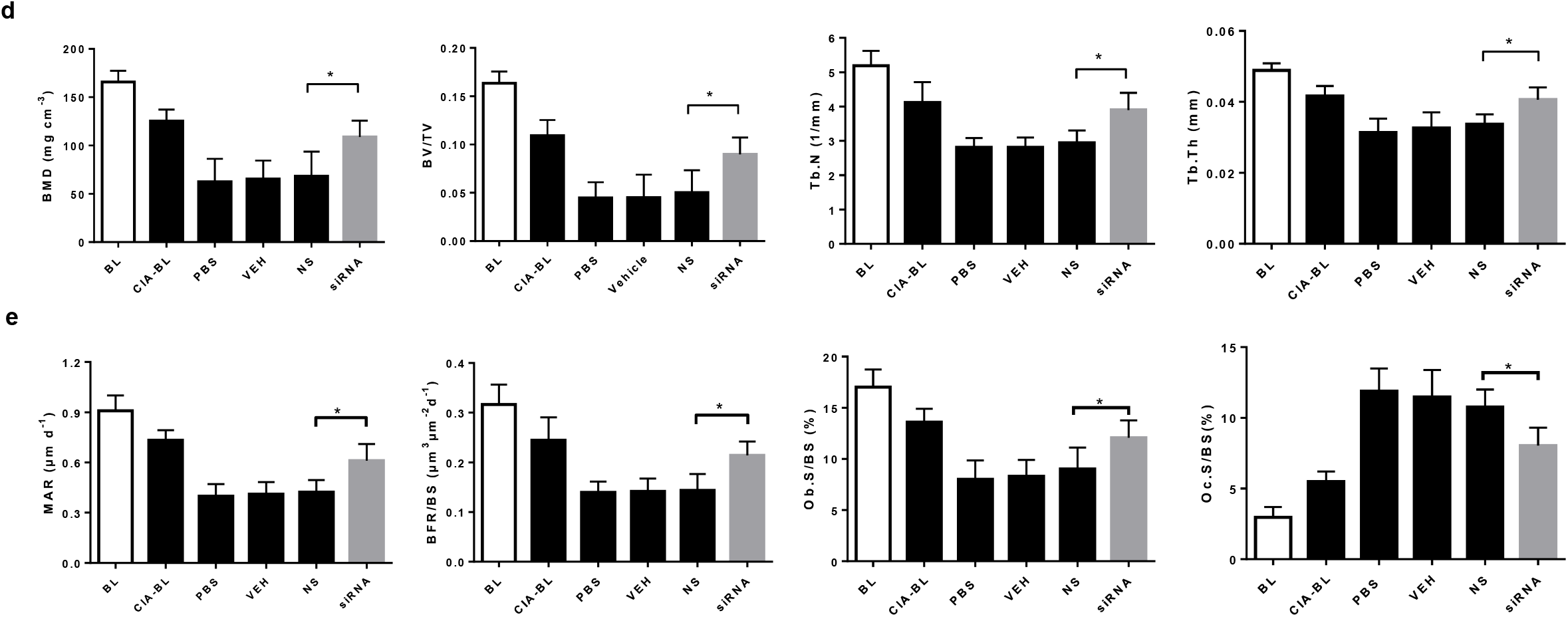
Osteoblastic PLEKHO1 inhibition leads to amelioration of articular inflammation and promotion of bone formation in CIA mice. (**a**) Time course changes in arthritis score from CIA mice treated with PBS, VEH, NS, and siRNA, respectively. (**b**) Histological inflammation score in the hind paws from the CIA mice after 6-week treatment in the respective group. (**c**) Representative histological images (Scale bars, 50 μm), microCT images (proximal tibial bones, Scale bars, 100 μm), calcein green and xylenol orange double labelling (Scale bars, 10 μm) from the hind paws of mice in each group at day 70 after primary immunization. (**d**) The values of microCT parameters (BMD, BV/TV, Tb.N and Tb.Th) of the proximal tibial bones from the CIA mice after treatment in the respective group. (**e**) The values of bone histomorphometric parameters (MAR, BFR/BS, Ob.S/BS and Oc.S/BS) in the hind paws from the CIA mice after treatment in the respective group. All data are the mean ± s.d. *P < 0.05, **P < 0.01. *n* = 7 per group. For **a**, a two-way ANOVA with subsequent Bonferroni posttests was performed. For **b,d,e**, a one-way ANOVA with subsequent Tukey’s multiple comparisons test was performed. Note: BL, baseline; CIA-BL, collagen-induced arthritis baseline; VEH, (AspSerSer)_6_-liposome; NS, (AspSerSer)_6_-liposome -NS siRNA; siRNA, (AspSerSer)_6_-liposome -*Plekho1* siRNA.

### Plekho1 siRNA delivered to osteoblasts improves arthritis symptoms and promotes bone formation in a collagen-induced non-human primate arthritis model

To facilitate our findings to clinical translation, we carried out *Plekho1* siRNA inhibition experiments in cynomolgus monkeys (**Supplementary Fig. 8a**). We firstly confirmed that *Plekho1* mRNA expression in osteoblasts was significantly inhibited till 5 weeks after treatment with siRNA (**Supplementary Fig. 8b**). Thereafter, we found that the body weight, an important indicator for disease progression, in NS and PBS-treated monkeys were gradually decreased during 5-week treatment, whereas the body weight in siRNA-treated monkeys were continuously increased except one monkey (Monkey 8) (**Fig. 6a**). We also found that arthritis score and the proximal interphalangeal point (PIP) joint swelling were continuously increased in NS and PBS-treated monkeys, whereas these two parameters in siRNA-treated monkeys maintained at a low level during 5-week treatment except Monkey 8 (**Fig. 6b, c**). We performed histological evaluation and found that joint inflammation in siRNA-treated monkeys was significantly attenuated than that of NS and PBS-treated monkeys after treatment (**Fig. 6d, 6e**). We then used X ray to assess the bony structure of the PIP joints and found that siRNA treatment could alleviate the continuous increase of the PIP radiological score. After treatment, the average PIP radiological score in PBS, NS and siRNA-treated monkeys were 43.33 ± 5.51, 43.67±7.09, 20±14.42, respectively (**Fig. 6f, 6g**). Consistent with the X ray findings, microCT reconstruction images also showed better organized bony and joint structure in siRNA-treated monkeys compared to NS and PBS-treated monkeys after treatment (**Fig. 6h**). The values of BMD and BV/TV were significantly increased in siRNA-treated monkeys after treatment, whereas such changes were not observed in monkeys treated with NS and PBS (**Fig. 6i**). Furthermore, we found a larger width between the two labeling bands in the siRNA-treated monkeys compared to NS and PBS-treated monkeys after treatment (**Fig. 6j**). The values of bone formation parameters MAR and BFR/BS were also significantly higher in siRNA-treated monkeys compared to NS and PBS-treated monkeys after treatment (**Fig. 6k**). In addition, we found that siRNA treatment did not cause any visible differences in appearance, behavior or sign of organ damage when examined by necropsy. To further evaluate its safety, we performed routine blood, blood coagulation and blood biochemistry tests, and found no statistically significant differences in these parameters among the three groups during 5-week treatment (**Supplementary Fig. 8c-e**).

**Figure 6.**
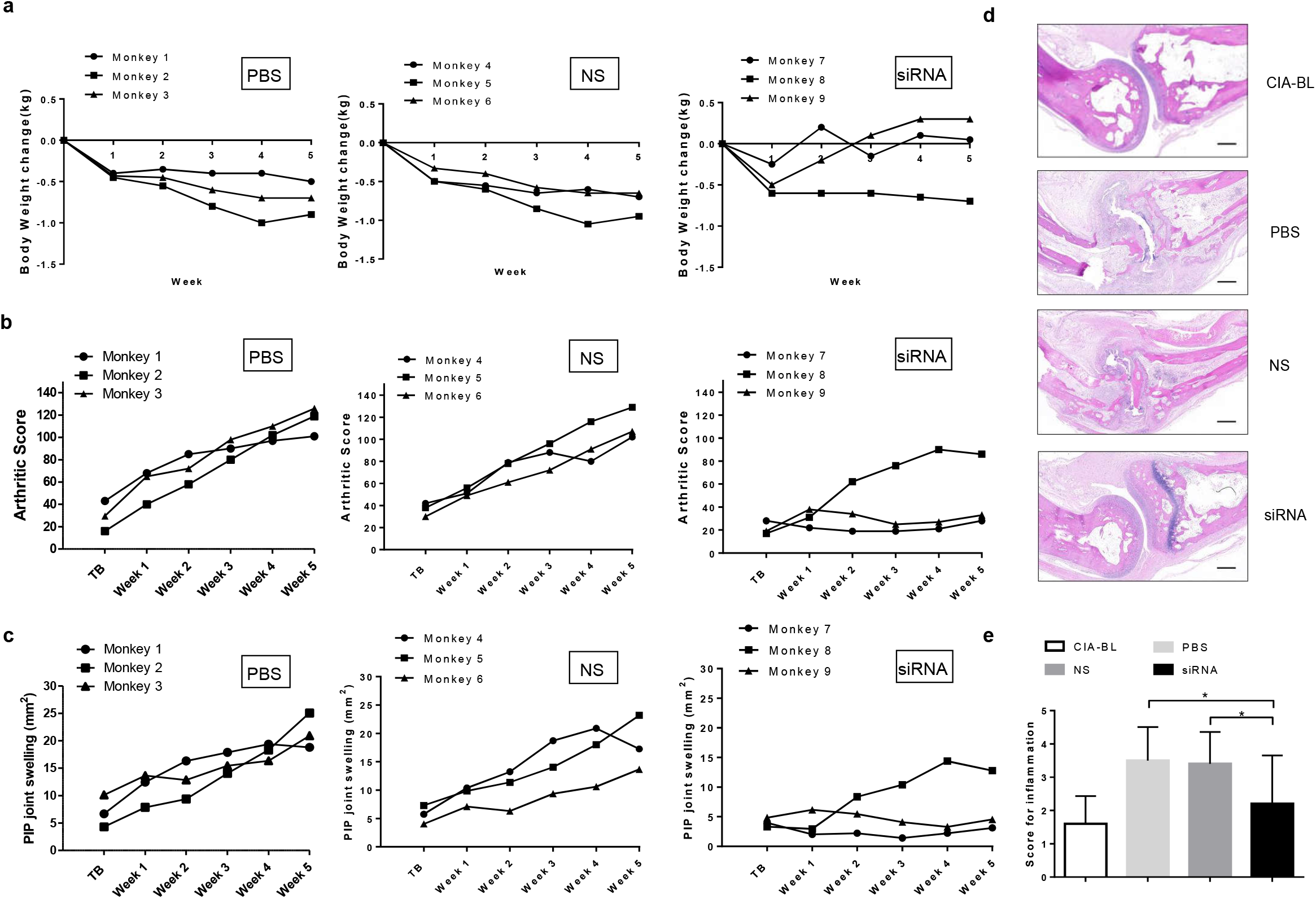
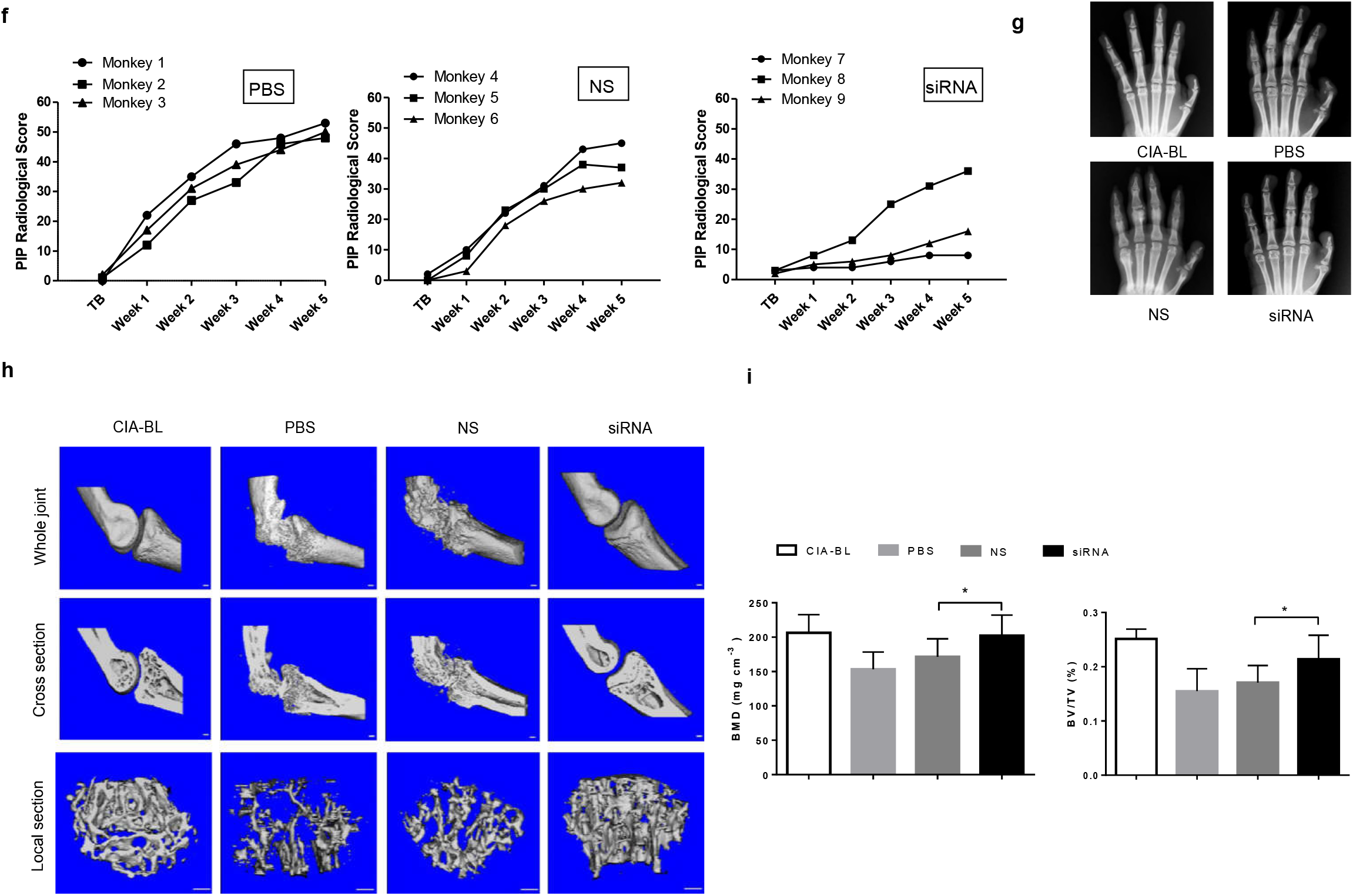
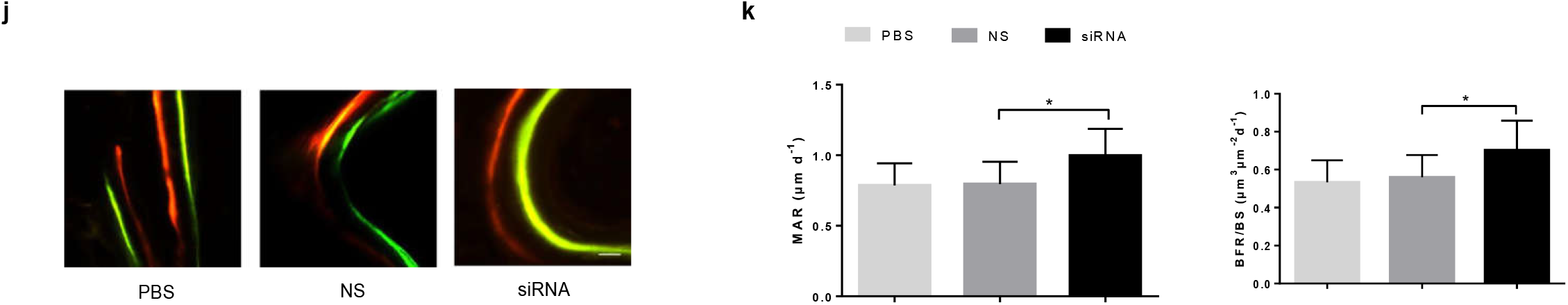
RNA interference-based silencing of PLEKHO1 with osteoblast-selective delivery rescues articular inflammation and bone erosion in non-human primate arthritis model induced by type II bovine collagen. (**a**) Body weight monitoring over time of each cynomolgus monkey after treatment. (**b**) Analysis of arthritis score over time of each cynomolgus monkey after treatment. (**c**) PIP joints swelling monitoring over time of each cynomolgus monkey after treatment. (**d**) Representative histological change of PIP joints from the hand of cynomolgus monkey in each group (four PIPs in each cynomolgus monkey, three cynomolgus monkeys in each group). Scale bar, 500 μm. (**e**) Comparison of histological inflammation score in each group. (**f**) Radiological score monitoring over time by X ray examination of the hand in each cynomolgus monkey. (**g**) Representative X ray image of the hand in each group (three cynomolgus monkeys in each group). (**h**) Representative 3D microarchitecture of the PIP in each group (four PIPs in each cynomolgus monkey, three cynomolgus monkeys in each group), obtained by microCT examination. Scale bars, 1.0 mm. (**i**) Analysis of the three-dimensional architecture parameters (BMD and BV/TV) for monkeys in each group. (**j**) Bone formation was examined by sequential labels with fluorescent dye in nondecalcified bone sections from cynomolgus monkeys. Representative fluorescent micrographs of PIP showed the xylenol (red) and calcein (green) labels in each group (four PIPs in each cynomolgus monkey, three cynomolgus monkeys in each group). Scale bar, 10 μm. (**j**) Analysis of bone histomorphometric parameters (MAR and BFR/BS) from cynomolgus monkeys in each group. All data are the mean ± s.d. *P<0.05. A one-way ANOVA with subsequent Tukey’s multiple comparisons test was performed. Note: CIA-BL, collagen-induced arthritis baseline; NS, (AspSerSer)_6_-liposome -NS siRNA; siRNA, (AspSerSer)_6_-liposome *-Plekho1* siRNA; TB, treatment begins; PIP, proximal interphalangeal.

## Discussion

This study, for the first time, uncovered a new role of osteoblastic PLEKHO1 in regulating joint inflammation in RA pathogenesis. We further demonstrated that osteoblastic PLEKHO1 inhibition could not only augment bone formation but also inhibit joint inflammation in RA.

Previous views suggest that the joint inflammation in RA is mainly mediated by the activated inflammatory cells such as FLS and immune cells including infiltrated T lymphocytes and leukocytes[20]. However, it remains unclear whether other cell populations also contribute to regulate the local joint inflammation. Albeit osteoblasts are among the dominant cell types in the joint, the current perspectives restrict their role in the modulation of bone formation in RA. Interestingly, our study showed that osteoblasts could directly contribute to regulate the local joint inflammation during RA development. The inflammation environment of RA could induce the expression of PLEKHO1 within osteoblasts, as evidenced by the aberrant high levels of PLEKHO1 in osteoblasts of the joint bone samples from both RA patients and CIA mice as well as the increased PLEKHO1 expression in both human and mouse osteoblast-like cells stimulated by various inflammatory cytokines. More surprisingly, we found that genetic deletion of Plekho1 gene in osteoblasts markedly attenuated the local joint inflammation in inflammatory arthritic mouse models, as supported by the low arthritis scores and histological inflammation scores as well as downregulated inflammatory cytokines (IL-1β and IL-6) in joint samples from the *Plekho1_osx_*^-/-^ mice induced by type II collagen or K/BxN serum. Moreover, conditional overexpression of osteoblastic Plekho1 in the Plekho1 systemic knockout mice (*Plekho1*^-/-^/*Plekho1_osx_ Tg*) significantly exacerbated the joint inflammation in CIA model and STA model. All these findings indicate that the osteoblasts with high PLEKHO1 expression contribute to the joint inflammation in RA. Despite the regulatory role on BMP signaling, PLEKHO1 has been identified to regulate the cytokine signaling response to inflammation[10, 11]. A recent study showed that overexpression of PLEKHO1 triggered the activation of pro-inflammatory pathways in the HEK293 model system[21]. Consistently, we found lower levels of inflammatory cytokines (IL-1β and IL-6) in CD4+T cells and FLS as well as decreased migration of neutrophils after co-culture with the *Plekho1*-depleted osteoblasts derived from the *Plekho1_osx_*^-/-^ mice when compared to those co-cultured with Plekho1-intact osteoblasts. Furthermore, the above changes in the inflammatory cells were reversed after co-culture with the *Plekho1*-overexpressed osteoblasts derived from the *Plekho1_osx_ Tg* mice. All these results suggest that the elevated osteoblastic PLEKHO1 could regulate the inflammatory cells that were commonly activated in the local inflammatory environment in RA joint.

TNF-α plays a pivotal role in regulating the inflammatory response in RA through inducing profinflammatory cytokines release, activating inflammatory cells and promoting leukocytes accumulation[22]. Recently, emerging evidence indicated that osteoblasts could respond to the stimulations of TNF-α by secretion of several inflammatory effectors[23, 24]. Our studies showed that silence of osteoblastic Plekho1 could remarkably decrease the production of the important pro-inflammatory cytokines (IL-1β and IL-6). Moreover, we observed remarkable alleviation of local inflammation in human TNF-α transgenic mice with osteoblastic deficiency of Plekho1, implying that osteoblastic PLEKHO1 on joint inflammation regulation might be related with TNF-α involved signal mechanism. Therefore, to further investigate the molecular mechanism of osteoblastic PLEKHO1 modulating the joint inflammation in RA, we performed a series of *in vitro* studies. We found that the phosphorylation of IKKα/β and p65 as well as the ubiquitination of RIP1 in NF-κB signal pathway were remarkably inhibited in TNF-α-stimulated osteoblasts after PLEKHO1 silence. Moreover, we confirmed that PLEKHO1 could bind to TRAF2 and promote TRAF2-mediated RIP1 ubiquitination in TNF-α-stimulated osteoblasts, and the domains for the interaction were LZ-containing domain of PLEKHO1 and C domain of TRAF2 respectively. In contrast to the previous studies which demonstrated that lipid mediator sphingosine-1-phosphate specifically bound to TRAF2 and increased TRAF2-catalyzed ubiquitination of RIP1[25], our study found a new cofactor “PLEKHO1” required for TRAF2-mediated RIP1 ubiquitination to activate NF-kB for inducing inflammatory cytokines production (**Supplementary Fig. 9**).

In addition, we observed that the osteoblast-specific *Plekho1*-depleted CIA mice showed an apparent attenuation in bone formation reduction and osteoblast-targeted Plekho1 siRNA treatment could promote bone repair in CIA rodents, which could be explained by the above documented role of PLEKHO1 as a critical ubiquitylation modulator for regulating the BMP signaling[7]. Certainly, PLEKHO1’s effect on local inflammation regulation might also be considered as one of the mechanisms for its bone formation modulation, as the common view regarded that inflammation could impair the function of osteoblasts-mediated bone formation in RA[26].

Considering the role of osteoblastic PLEKHO1 in regulating joint inflammation and bone erosion repair during RA, inhibition of it may exert dual beneficial effect on RA treatment. Thus, to facilitate its clinical translation, we evaluated the effect of osteoblast-targeted PLEKHO1 inhibition by administrating *Plekho1* siRNA encapsulated by our developed osteoblast-targeting delivery system in a CIA mouse model and a non-human primate arthritis model. We demonstrated that osteoblast-targeting *Plekho1* siRNA treatment not only improved arthritis symptoms and but also promoted bone formation in both animal models. Moreover, the siRNA treatment didn’t cause any significant side effects. These results will also facilitate clinical translation of RNA interference-based therapeutic strategy in RA, since currently reports about siRNA-based therapeutic approaches for rheumatic diseases are almost from rodent experiments[27].

In summary, our studies uncovered a previously unrecognized role of osteoblastic PLEKHO1 in mediating the joint inflammation during RA development. Targeting osteoblastic PLEKHO1 may be a promising therapeutic strategy with dual action of inflammation inhibition and bone formation augmentation for RA.

## Methods

### Human joint bone tissue preparation

Tibial plateau samples from knee joint of subjects with RA (fulfilled the 1987 revised American College of Rheumatology criteria) were obtained at knee joint replacement surgery and provided by Shanghai Guanghua Hospital (China). Subjects with malignancy, diabetes or other severe diseases in the previous 5 years were excluded from our study. For non-RA control, we selected age- and sender-matched subjects who diagnosed as severe trauma (TM) and underwent knee joint replacement surgery in Shanghai Guanghua Hospital and Shenzhen 8th People Hospital (China). Half of the bone samples were fixed with 4% buffered formalin before decalcification, the other half were stored in liquid nitrogen until use. All the clinical procedures were approved by the Committees of Clinical Ethics in the Shanghai Guanghua Hospital and Shenzhen 8th People Hospital. All subjects gave informed consent prior to surgery.

### Cells

Murine osteoblastic cell line, MC3T3E1, was purchased from the Cell Resource Center, Peking Union Medical College (China) and cultured in α-MEM (Thermo Fisher Scientific) containing 10% fetal bovine serum (FBS, Thermo Fisher Scientific) at 37°C in a humidified 5% CO2 incubator. Human osteoblast–like cell line, MG-63, was purchased from ATCC and cultured in EMEM (Thermo Fisher Scientific) containing 10% FBS at 37°C in a humidified 5% CO2 incubator. Human embryonic kidney HEK293T cell was purchased from ATCC and cultured in DMEM (Thermo Fisher Scientific) containing 10% FBS at 37°C in a humidified 5% CO2 incubator. Primary osteoblast precursor cells were isolated from the calvarial bone of newborn C57BL/6 mice (1- to 2-day-old) through enzymatic digestion with α-MEM containing 0.1% collagenase (Life technologies, Grand Island, NY, USA) and 0.2% dispase II (Life Technologies, Grand Island, NY, USA). The isolated osteoblast precursor cells were promoted with osteogenic α-MEM medium with 10% FBS, 1% penicillin-streptomycin, 5mM β-glycerol phosphate (Sigma-Aldrich, St. Louis, MO, USA), 0.1mg/ml ascorbic acid (Sigma-Aldrich, St. Louis, MO, USA) and 10nM dexamethasone (Sigma-Aldrich, St. Louis, MO, USA) for 9 days and culture medium was replaced every 2–3 days[28]. Primary fibroblast-like synoviocytes (FLS) were isolated from the ankle joints of C57BL/6 mice by collagenase digestion[29]. Isolated FLS were grown in DMEM high glucose, GlutaMAX (Life Technologies, Grand Island, NY, USA) supplemented with 10% FBS and 1% penicillin-streptomycin. As no bone marrow-derived lineage cells were detected after passage 3, cultured FLS were used between passages 4-8. CD4+T lymphocytes were isolated from the spleen single-cell suspension of C57BL/6 mice by specific magnetic beads-based negative-selection (Miltenyi Biotec, Bergisch Gladbach, Germany). Neutrophils were isolated from peripheral blood of C57BL/6 mice by specific magnetic beads sorting (Miltenyi Biotec, Bergisch Gladbach, Germany)[30]. Before use, osteoblasts were confirmed to express osteocalcin, synovial fibroblasts were confirmed to express vascular cell adhesion molecule 1 and not F4/80 or CD45, CD4+T lymphocytes were confirmed to express CD3 and CD4, and neutrophils were confirmed to express Gr-1 and CD11b.

### Co-culture and migration experiment

For osteoblasts-FLS co-culture: FLS were grown in co-culture with osteoblasts from *Plekho1_osx_*^-/-^ mice, *Plekho1_osx_ Tg* mice or WT littermates. Co-culture experiments were conducted in 6-well transwell coculture plates (Millipore, Billerica, MA, USA). The primary osteoblasts (5×10^5^/well) were placed in the lower well and FLS (5×10^5^/well) in the inserts. TNF-α was added for 24 h. After experiment, FLS were collected and the mRNA levels of IL-1 β and IL-6 were detected by quantitative real-time PCR. For osteoblasts-CD4+T cells co-culture: Isolated CD4+T lymphocytes were grown in co-culture with osteoblasts from *Plekho1_osx_*^-/-^ mice, *Plekho1_osx_ Tg* mice or WT littermates. Coculture experiments were conducted in 6-well transwell coculture plates (Millipore, Billerica, MA, USA). The primary osteoblasts (5×10^5^/well) were placed in the lower well and CD4+T lymphocytes (1×10^6^/well) in the inserts. TNF-α was added for 24 h. After experiment, FLS were collected and the mRNA levels of IL-1β and IL-6 were detected by quantitative real-time PCR. For osteoblasts-neutrophils co-culture: Isolated neutrophils were plated onto 3.0μm pore polycarbonate membrane transwell inserts, which were placed into the osteoblasts culture wells in the presence of TNF-α. Neutrophils were allowed to migrate through the membrane for 2 h. Migrated cells attached to the bottom of the well were stained by chloroacetate esterase and positive cells were counted to determine changes in chemotactic activity of osteoblasts.

### siRNA inhibition

Plekho1 siRNA and non-sense siRNA (random siRNA) were purchased from GenePharma (Shanghai, China). Transfection was performed with Lipofectamine^™^ 2000 (Invitrogen Life Technologies, Carlsbad, CA, USA) according to the manufacturer’s instructions. Briefly, murine osteoblastic cell line MC3T3E1 were seeded in a 6-well plate. Then, 4 ul of 20 uM Plekho1 siRNA and 2.5 ul of Lipofectamine^™^ 2000 were diluted in 0.25 ml of Opti-MEM separately for each well. After 5 min of incubation at room temperature, the diluted Plekho1 siRNA and Lipofectamine^™^ 2000 were mixed gently and allowed to stand for 20 min at room temperature. Then, cells were washed by PBS. And 1.5 ml MEM without serum and antibiotics was added to each well. Followed by adding 0.5 ml of Opti-MEM containing the Lipofectamine^™^ 2000-siRNA complex and incubated for 6 h at 37 °C. Non-sense siRNA transfected by Lipofectamine^™^ 2000 were used as negative control. Then, the Lipofectamine^™^ 2000-siRNA complex was removed and replaced with fresh complete MEM. Next, the cells were stimulated by 10 ng/ml TNF-α for another 45 min. The supernatant and the cell lysis were collected for ELISA and western blot analysis, respectively.

### Plasmid construction and transfection

HEK293T cells were cultured and used for transfection. Plasmids of different epitope-tagged human Plekho1 and TRAF2, including both full-length proteins and truncated mutants, as well as human RIP1 were constructed by PCR, followed by subcloning into expression vector pcDNA3.1[17, 18, 31, 32]. Plasmid of HA-tagged human ubiquitin-K63 was from Addgene. Transfection was performed using Lipofectamine 2000 (Invitrogen) according to the manufacturer’s instructions [7].

### Immunoprecipitation and immunoblotting

At 36 h after the transfection, HEK293T were lysed in HEPES lysis buffer (20 mM HEPES pH 7.2, 50 mM NaCl, 0.5% Triton X-100, 1 mM NaF, 1 mM dithiothreitol) supplemented with protease inhibitor cocktail (Roche) and phosphatase inhibitors (10mM NaF and 1mM Na_3_VO_4_). Immunoprecipitations were performed using anti-Flag or anti-Myc antibody and protein A/G-agarose (Santa Cruz Biotechnology) at 4 °C. Plekho1 and TRAF2, including full-length proteins and truncated mutants, in lysates or immunoprecipitates were examined by anti-Myc (Cell Signaling Technology) and anti-Flag (Sigma) primary antibodies, respectively, and the appropriate secondary antibodies in immunoblotting, followed by detection with SuperSignal chemiluminescence kit (Thermo Fisher Scientific)[6, 7, 33].

### *In vivo* ubiquitylation assay

HEK293T cells were transfected with Myc-RIP1, HA-Ub-K63, Flag-Plekho1 or Flag-ΔLZ and Flag-TRAF2 or Flag-ΔTRAF C. At 36 h after the transfection, cells were lysed in HEPES lysis buffer and then incubated with anti-Myc antibody (Sigma) for 3 h and protein A/G-agarose beads (Santa Cruz Biotechnology) for a further 8 h at 4 °C. After three washes, ubiquitination of RIP1 was detected by immunoblotting with an anti-HA antibody (Cell Signaling Technology)[7, 25].

### Animals

Osteoblast-specific *Plekho1* knockout mice *(Plekho1_osx_*^-/-^ mice), *Plekho1* knockout mice *(Plekho1*^-/-^ mice) and osteoblast-specific *Plekho1* overexpressing *(Plekho1_osx_ Tg*) mice were prepared according to our previous reports[34]. We generated the *Plekho1_osx_*^-/-^ mice based on the Cre-loxP strategy. Briefly, we created the heterozygous mice carrying the mutant allele with LoxP sites harboring exon 3 to exon 6 of Plekho1 gene *(Plekho1^fl/-^),* which were then crossed with Osx-Cre mice (Beijing Biocytogen Co., Ltd, China) to generate the *Plekho1_osx_*^-/-^mice. To generate *Plekho1*^-/-^ mice, *Plekho1*^*fl*/+^ mice were crossed with CMV-Cre mice (Beijing Biocytogen Co., Ltd, China). To generate *Plekho1_osx_ Tg* mice, we created a mouse strain carrying the ROSA26-PCAG-STOP^fl^-Plekho1-eGFP allele, and crossed them with the Osx-Cre mice to generate the *Plekho1_osx_ Tg* mice that overexpressing Plekho1 in osteoblasts. DBA/1 mice and C57BL/6J mice were from Vital River Laboratory Animal Technology Co., Ltd. (Beijing, China). The human TNF transgenic (hTNFtg) mice (strain Tg197; genetic background, C57BL/6) were obtained from Dr. G. Kollias (Institute of Immunology, Biomedical Sciences Research Center “Alexander Fleming,” Vari, Greece). KRN T cell receptor (TCR) transgenic mice (B10.BR genetic background) were obtained from Dr. D. Mathis and Dr. C. Benoist (Harvard Medical School, Boston, Massachusetts, USA) and maintained on a C57BL/6 background. NOD/Lt mice were purchased from HFK Bioscience Co.,Ltd (Beijing, China) and were used for breeding of K/BxN arthritic mice. Male *Plekho1_osx_*^-/-^ mice were bred with female hTNFtg mice to obtain *Plekho1_osx_*^-/-^/hTNFtg mice. Male *Plekho1*^-/-^ mice were bred with female *Plekho1_osx_ Tg* mice to generate offspring *(Plekho1^-/-^/Plekho1_osx_ Tg* mice) that express high PLEKHO1 exclusively in osteoblasts. All mice data were generated from age- and sex-matched littermates. Female cynomolgus monkeys (Macaca fascicularis), 3-4 years old, were purchased from PharmaLegacy Laboratories Co., Ltd (Shanghai, China). All the animals were maintained under standard animal housing conditions (12-h light, 12-h dark cycles and free access to food and water). All procedures were approved by the Ethical Animal Care and Use Committee in China Academy of Chinese Medical Sciences and Hong Kong Baptist University.

### Collagen-induced arthritis (CIA) mice model

The induction of CIA was performed as previously described with slight modification[35]. Briefly, type II collagen (bovine type II collagen for DBA/1 mice, chicken type II collagen for C57BL/6J mice) (Sigma-Aldrich, St. Louis, MO, USA) was dissolved at a concentration of 2 mg/ml in 0.05 M acetic acid and emulsified with complete Freund’s adjuvant (CFA) (Sigma-Aldrich, St. Louis, MO, USA). To induce CIA in C57BL/6J mice, 100mg of desiccated killed *Mycobacterium tuberculosis* H37Ra (BD Biosciences) was suspended in 30ML of CFA[36]. On day 0, the CIA mice were immunized with 0.1 mL of collagen by intradermal injection at the base of the tail. On day 21, mice were given a booster dose of collagen through the same route.

### K/BxN serum-transfer arthritis (STA) mice model

Pro-arthritic serum was isolated from K/BxN mice (generated by crossing KRN TCR transgenic mice with NOD/Lt mice) as previously described[37]. K/BxN arthritis was induced by the intraperitoneal injection of diluted K/BxN serum (75 μl serum with 75 μl endotoxin-free PBS) in C57BL/6J mice on day 0 and 2[38].

### Collagen-induced arthritis (CIA) non-human primate model

The induction of CIA in cynomolgus monkeys was performed as previously described with slight modification[39]. Briefly, bovine type II collagen (4 mg/mL) was emulsified in an equal volume of complete Freund’s adjuvant. Ten monkeys were immunized with 1 mL of the cold emulsion by intradermal injections divided over 20 sites: 19 sites on the back and 1 site at the base of the tail, and after a three-week interval, boosted with the same immunizing procedure. During the study, body weight was recorded once a week. Onset of CIA was monitored by arthritic score, once a monkey fulfilled the criteria of 5% of the maximum arthritic score, it was assigned to the respective treatment group.

### Treatment

For mice siRNA treatment experiment: DBA/1 mice were induced by type II collagen. After CIA was successfully established (day 28 after primary immunization, a mouse with a score of one or above was regarded as arthritic), parts of the mice were sacrificed before treatment as CIA baseline (CIA-BL). The remaining CIA mice were divided into (AspSerSer)_6_-liposome-Plekho1 siRNA group, (AspSerSer)_6_-liposome-non sense siRNA group, (AspSerSer)_6_-liposome group, and vehicle control group. The mice in each group received six periodic intravenous injections of (AspSerSer)_6_-liposome-*Plekho1* siRNA, (AspSerSer)_6_-liposome-non sense siRNA, (AspSerSer)_6_-liposome, and PBS, respectively, with a siRNA dose of 5.89 mg/kg at an interval of one week.

For non-human primate siRNA treatment experiment: After CIA was successfully established, one monkey was sacrificed before siRNA treatment as CIA baseline (CIA-BL). The remaining CIA monkeys were divided into three groups: (AspSerSer)_6_-liposome-*Plekho1* siRNA group, (AspSerSer)_6_-liposome-non sense siRNA group, and vehicle control group. The monkeys in each group received five periodic intravenous injections of (AspSerSer)_6_-liposome-*Plekho1* siRNA, (AspSerSer)_6_-liposome-non sense siRNA and PBS, respectively, with a siRNA dose of 1mg/kg at an interval of one week.

### Evaluation of arthritis severity

Clinical severity of arthritis in mice was evaluated according to the following visual scoring system[35]: 0 = no swelling or erythema; 1 =Erythema and mild swelling confined to the tarsals or ankle joint; 2 = Erythema and mild swelling extending from the ankle to the tarsals; 3 = Erythema and moderate swelling extending from the ankle to metatarsal joints; 4 = Erythema and severe swelling encompass the ankle, foot and digits, or ankylosis of the limb. Each limb was assigned a score of 0 to 4, with a maximum possible score of 16 for each mouse.

The severity of arthritis in monkeys was assessed by observing the degree of swelling and rigidity at the metacarpophalangeal (MCP), proximal interphalangeal (PIP), and distal interphalangeal (DIP) joints of both the hands and feet, as well as the wrists, ankle, elbow, and knee (total 64 joints) weekly until the end of experiment, as reported in previous literature with slight modifications[40]. Each joint was assessed in accordance with the evaluation criteria shown as follows: 0 = normal; 1 = swelling not visible but can be determined by touch; 2 = swelling slightly visible and can be confirmed by touch; 3 = swelling clearly visible and/or noticeable joint deformity. Each joint is examined as per the evaluation criteria. The arthritis score for each animal was designated as the sum of individual joint score, with maximum score of 192.

### Evaluation of joint swelling

Joint swelling at PIP joints of each cynomolgus monkey was measured by vernier caliper (Mitutoyo, Neuss, Germany) every week and calculated using the following formula: PIP swelling(mm2)= horizontal diameter x vertical diameter x 3.14 x 0.25.

### Radiographic Examination

Radiographs of the animals were obtained using a Portable X-ray SP-VET-4.0 unit (Sedecal, Madrid, Spain). In anesthetized condition, the hands and feet of the monkeys were radiographed in the dorso-palmar position. Radiology projections of arthritic hands and feet were performed weekly to assess the extent of joint destruction, bone erosion and joint space narrowing. The digital radiographs images of the hands and feet were acquired and the PIP joints were graded according to the modified Larsen’s method with minor modifications[41]. The scoring criteria were as follows: 1 = Minor deformity in joint cartilage layers, and / or subchondral bone regions; 2 = Severe deformity in joint cartilage layers and subchondral bone regions, small amount of osteophytes present. The joint cavity fuzzy, but visible; 3 = Severe Grade 2 changes + Large amount of osteophytes present. The joint cavity was indistinguishable or invisible; 4= Severe Grade 3 changes + the joint cavity undetectable. Bones appeared sclerotic or ankyolosing with major joint disfiguration. The scoring was done by two blinded professional investigators respectively. The radiologic score of PIP joints (4 x4) for each animal ranged from 0 to 64.

### Routine blood tests and blood biochemistry

During the course of non-human primate studies, blood samples (2 mL) were collected in EDTA treated tubes at various time points and processed appropriately to run assessments for hematology, coagulation, serum chemistry. Hematology and coagulation measurements were made on whole blood and analyzed with automated hematology analyzer (Sysmex XT-2000iV^™^, UK) and Amax Destiny Plus^™^ analyzer (Trinity Biotech), respectively. Serum chemistry was determined with biochemical analyzer Hitachi-7180 (Hitachi, Tokyo, Japan).

### ELISA

Ankle joints of mice were pulverized using a mortar and pestle filled with liquid nitrogen. Tissue was transferred to 15 ml tubes, placed on dry ice and resuspended in 1 ml PBS/200 mg of tissue and homogenized using a tissue homogeniser. Joint homogenates were centrifuged for 10 min at 500g at 4°C. Supernatants were transferred to 1.5 ml tubes, centrifuged at 15,000 g for 5 min and collected for analysis. IL-1β and IL-6 quantifications were determined by ELISA according the instructions of the manufacturer (eBioscience), data was multiplied by a dilution factor for conversion from pg/ml to pg/g [42].

### Total RNA isolation, reverse transcription and quantitative real-time PCR

Total RNA was isolated from cultured cells and isolated cells using the RNeasy^®^ Mini Kit (QIAGEN, Dusseldorf, Germany). The concentration of total RNA was determined using a spectrophotometer. cDNA was synthesized from 0.5 μg of total RNA using a commercial first-strand cDNA synthesis kit (QIAGEN, Dusseldorf, Germany). Real-time PCR reactions were performed using SYBR Green in a 7900HT Fast Real-Time PCR System (Applied Biosystems, Foster City, California, USA). The employed primer sequences in the study were listed in **Supplementary Table 1**. The amplification conditions were as follows: 50°C for 2 minutes, 95°C for 10 minutes, 40 cycles of 95°C for 15 seconds, and 60°C for 1 minute. The fluorescence signal emitted was collected by ABI PRISM^®^ 7900HT Sequence Detection System and the signal was converted into numerical values by SDS 2.1 software (Applied Biosystems). The mRNA expression level of the target gene was first calculated from the Relative Standard Curve Method by the SDS 2.1 software. The threshold cycle (CT) value, which represents the relative expression of each target gene, was determined from the corresponding curve. Then, the relative expression of mRNA was determined by dividing the target amount by endogenous control amount to obtain a normalized target value. The relative mRNA expression was calculated using the 2^-ΔΔCt^ method as follow: ΔΔCt = (Ct_target gene_ – Ct_reference gene_) experimental group – (Ct_target gene_ Ct_reference gene_) _WT-OB group_)[8].

### Western blot analysis

The bone specimens and cells were snap-frozen in liquid nitrogen and mechanically homogenized in lysis buffer. Protein fractions were collected by centrifugation at 15,000g at 4 °C for 10 min and then subjected to SDS-PAGE and transferred to polyvinylidene difluoride (PVDF) membranes. The membranes were blocked with 5% BSA and incubated with specific antibodies overnight. A horseradish peroxidase–labeled secondary antibody (Abcam) was added and visualized using an enhanced chemiluminescence kit (Pierce). Antibodies to the following proteins were used: GAPDH (Sigma), PLEKHO1 (Santa Cruz, sc-50227), IKKβ (Cell Signaling Technology, #8943), p-IKKα/β (Cell Signaling Technology, #2078), p65 (Cell Signaling Technology) and p-p65 (Abcam, ab106129). The relative amounts of the transferred proteins were quantified by scanning the auto-radiographic films with a gel densitometer (Bio-Rad, USA) and normalized to the corresponding GAPDH level[43].

### Laser capture micro-dissection (LCM)

For LCM, the hind paw of mice and the hand of monkeys were decalcified in 10% ethylenediaminetetraacetic acid (EDTA) and embedded in OCT. Then, the series frozen sections (5μm) were prepared in a cryostat (CM3050; Leica Microsystems, Wetzlar, Germany) at -20°C. Adjacent sections were mounted on either glass slides or polyethylene membrane–equipped slides (P.A.L.M., Bernried, Germany). The sections mounted on glass slides were performed immunostaining to identify the osteocalcin-positive cells. Briefly, the cryosections were air dried at room temperature, fixed in ice-cold acetone for 10 min, permeabilized in 0.1% Triton X-100 at room temperature for 20 min, and blocked in 5% donkey serum in PBS. The sections were incubated overnight at 4 °C with rabbit polyclonal anti-osteocalcin antibody (1:50 dilution; Santa Cruz Biotechnology, Inc.). After three washes in PBS, the sections were incubated with Alexa Fluor 488-conjugated donkey anti-rabbit IgG (1:400 dilution; Invitrogen). Finally, the sections were mounted with medium containing (4’,6-diamidinole-2-phenolindole) DAPI (Vector laboratories) and examined under a fluorescence microscope to identify the osteocalcin-positive staining cells. The adjacent sections mounted on membrane-coated slides were stained with neutral red for 1 min at room temperature. After brief rinsing in water, the sections were air-dried. Osteocalcin-positive staining cells in adjacent sections were isolated by microdissection with an upgraded laser pressure catapulting microdissection system (P.A.L.M.) using a pulsed 355 nm diode laser in the Leica LMD 7000 Laser Microdissection System. About 100~200 identified cells were collected in reaction tube containing 5μl lysis buffer for RNA extraction and subsequent real-time PCR analysis following above mentioned protocols[43].

### Immunofluorescence staining

The bone specimens were fixed with 4% buffered formalin and embedded with O.C.T. after decalcification with 10% EDTA. The frozen sections (5 μm thickness) were cut in a freezing cryostat at -20°C. The sections were air dried at room temperature, fixed in ice-cold acetone for 10 min, permeablilized with 0.1% Triton X-100 at room temperature for 20 min, and blocked in 5% donkey serum in PBS. The sections were then incubated overnight at 4°C with the mixture of goat polyclonal osteocalcin antibody (1:50; Santa Cruz Biotechnology, Inc.), and rabbit polyclonal PLEKHO1 antibody (1:50; Santa Cruz Biotechnology, Inc.). Following three washes in PBS, the sections were incubated with the mixture of Alexa Fluor 488-conjugated donkey anti-goat antibody, and Alexa Fluor 568-conjugated donkey anti-rabbit antibody (1:300; Invitrogen) for 1 h. Negative control experiments were done by omitting the primary antibodies. The sections were mounted with the medium containing DAPI (Vector Laboratories) and then examined under a fluorescence microscope (DM6000B, Leica image analysis system)[43, 44].

### MicroCT analysis

MicroCT analyses were performed using a high-speed μCT (viva CT 40, Scanco Medical, Switzerland) at energy of 70 kVp and intensity of 114 μA with high resolution and voxel size of 15 μm of 10.5 μm (mouse), 19μm (monkey). In mouse studies, the hind limbs were scanned and the proximal tibia was selected for analysis. In non-human primate study, the fore- hand was scanned and the PIP joint was selected for analysis. 3D reconstruction analysis was performed with SCANCO software (SCANCO Medical, Switzerland). Standard image analysis procedures were used to determine trabecular and cortical parameters. The following parameters were determined: bone mineral density (BMD), trabecular fraction (BV/TV), trabecular number (Tb.N) and trabecular thickness (Tb.Th)[45].

### Histology evaluation

The hind paw of mice and the hand of monkeys were fixed in 4% formaldehyde, decalcified with EDTA and embedded in paraffin. The serial sections of the paw and hand were examined for inflammation using hematoxylin and eosin staining. Inflammation was scored according to the following criteria: 0 = normal; 1 = minimal infiltration of inflammatory cells in perijoint area; 2 = mild infiltration; 3 = moderate infiltration; and 4 = marked infiltration[46]. Scoring was executed blindly by two investigators and mean values were calculated.

### Bone histomorphometric analysis

To assess bone formation, the animals were administered with xylenol orange (90 mg/kg) and calcein green (10 mg/kg) for labeling bone formation surface on day 12 and day 2 before sacrifice by intraperitoneal injection, respectively. The hind paw of mice and the hand of monkeys were fixed in 70% ethanol. The samples were trimmed from surrounding soft tissues, dehydrated in a series of ethanol solutions (70%, 80%, 90%, 95% and 100%), and embedded in methyl methacrylate. Non-decalcified histological sections (10μm thick) were made using a diamond saw microtome. The sections for light microscopic observations were performed by the modified Goldner’s trichrome, while unstained sections were prepared for fluorescent microscope observations. In mice study, the calcaneus area within the hind paw was chosen for histomorphometric analysis. In non-human primate study, the PIP joint was selected for analysis. All the measurements were performed using the professional image software (ImageJ, NIH, USA and BIOQUANT OSTEO, Version 13.2.6, Nashville, TN, USA). The following bone histomorphometric parameters for bone remodeling activity were analyzed, including mineral apposition rate (MAR), bone formation rate/bone surface (BFR/BS), osteoblast surface (Ob.S/BS), osteoblast number (Ob.N/B.Pm), osteoclast surface (Oc.S/BS) and osteoclast number (Oc.N/B.Pm)[8].

### Statistical analysis

For statistical analysis, data were expressed as the mean ± standard deviation. In general, statistical differences among groups were analyzed by one-way analysis of variance (ANOVA) with a Tukey’s multiple comparisons test to determine group differences in the study parameters. Especially, the dynamic detection data were compared by means of two-way ANOVA with subsequent Bonferroni posttests. All statistical analyses were performed using Graphpad Prism Software, 6.05 (Graph Pad Software Inc., San Diego, CA). P values less than 0.05 are considered significant.

## Acknowledgments

We thank the technical staff from Institute for Advancing Translational Medicine in Bone & Joint Diseases, Hong Kong Baptist University and Institute of Basic Theory, China Academy of Chinese Medical Sciences for providing critical comments and technical support. This study was supported by the Hong Kong General Research Fund (HKBU479111 to G.Z., HKBU478312 to G.Z., HKBU 12114416 to G.Z., HKBU262913 to G.Z., HKBU261113 to A.L., HKBU12122516 to A.L.), the Natural Science Foundation Council of China (81272045 to G.Z. and 81470072 to X.H.), the Research Grants Council & Natural Science Foundation Council of China (N_HKBU435/12 to G.Z.), the Hong Kong Baptist University Strategic Development Fund (SDF) (SDF15-0324-P02 to A.L.).

## Author Contributions

G.Z. and A.L. supervised the whole project. X.H., J.L., C.L., S.A.B., K.Z. and L.D. performed the major research and wrote the manuscript in equal contribution. B.G., D.L., C.L., Q.G. and D.F. provided the technical support. Y.B., H.F., L.X. and X.P. collected human bone specimens. C.X. and B.Z. provided their professional expertise.

